# SARS-CoV-2 ORF8 encoded protein contains a histone mimic, disrupts chromatin regulation, and enhances replication

**DOI:** 10.1101/2021.11.10.468057

**Authors:** John Kee, Samuel Thudium, David Renner, Karl Glastad, Katherine Palozola, Zhen Zhang, Yize Li, Joseph Cesare, Yemin Lan, Rachel Truitt, Fabian L. Cardenas-Diaz, Darrell N. Kotton, Konstantinos D. Alysandratos, Xianwen Zhang, Xuping Xie, Pei-Yong Shi, Wenli Yang, Edward Morrisey, Benjamin A. Garcia, Shelley L. Berger, Susan R. Weiss, Erica Korb

**Affiliations:** Department of Genetics, Perelman School of Medicine at the University of Pennsylvania, Philadelphia, PA 19104, USA; Department of Cell and Developmental Biology, Perelman School of Medicine at the University of Pennsylvania, Philadelphia, PA 19104, USA; Department of Microbiology, Perelman School of Medicine at the University of Pennsylvania, Philadelphia, PA 19104, USA; Department of Medicine, Perelman School of Medicine at the University of Pennsylvania, Philadelphia, PA 19104, USA; Department of Biochemistry and Biophysics, Perelman School of Medicine at the University of Pennsylvania, Philadelphia, PA 19104, USA; Department of Biology, Perelman School of Medicine at the University of Pennsylvania, Philadelphia, PA 19104, USA; Department of Epigenetics Institute, Perelman School of Medicine at the University of Pennsylvania, Philadelphia, PA 19104, USA; Department of Penn-CHOP Lung Biology Institute, Perelman School of Medicine at the University of Pennsylvania, Philadelphia, PA 19104, USA; Department of Penn Center for Research on Coronaviruses and Other Emerging Pathogens, Perelman School of Medicine at the University of Pennsylvania, Philadelphia, PA 19104, USA; Center for Regenerative Medicine, Boston University and Boston Medical Center and the Pulmonary Center and Department of Medicine, Boston University School of Medicine, Boston, MA 02118, USA; Department of Biochemistry and Molecular Biology, University of Texas Medical Branch, Galveston, TX, USA

## Abstract

SARS-CoV-2 emerged in China at the end of 2019 and caused the global pandemic of COVID-19, a disease with high morbidity and mortality. While our understanding of this new virus is rapidly increasing, gaps remain in our understanding of how SARS-CoV-2 can effectively suppress host cell antiviral responses. Recent work on other viruses has demonstrated a novel mechanism through which viral proteins can mimic critical regions of human histone proteins. Histone proteins are responsible for governing genome accessibility and their precise regulation is critical for a cell’s ability to control transcription and respond to viral threats. Here, we show that the protein encoded by ORF8 (Orf8) in SARS-CoV-2 functions as a histone mimic of the ARKS motif in histone 3. Orf8 is associated with chromatin, binds to numerous histone-associated proteins, and is itself acetylated within the histone mimic site. Orf8 expression in cells disrupts multiple critical histone post-translational modifications (PTMs) including H3K9ac, H3K9me3, and H3K27me3 and promotes chromatin compaction while Orf8 lacking the histone mimic motif does not. Further, SARS-CoV-2 infection in human cell lines and postmortem patient lung tissue cause these same disruptions to chromatin. However, deletion of the Orf8 gene from SARS-CoV-2 largely blocks its ability to disrupt host-cell chromatin indicating that Orf8 is responsible for these effects. Finally, deletion of the ORF8 gene affects the host-cell transcriptional response to SARS-CoV-2 infection in multiple cell types and decreases the replication of SARS-CoV-2 in human induced pluripotent stem cell-derived lung alveolar type 2 (iAT2) pulmonary cells. These findings demonstrate a novel function for the poorly understood ORF8-encoded protein and a mechanism through which SARS-CoV-2 disrupts host cell epigenetic regulation. Finally, this work provides a molecular basis for the finding that SARS-CoV-2 lacking ORF8 is associated with decreased severity of COVID-19.

## Main

SARS-CoV-2 has proven to be a highly virulent virus resulting in a devastating and global pandemic. Recent findings indicate that several other highly virulent viruses disrupt host cell epigenetic regulation through mimicry of host cell proteins^1–3^, particularly of histones^4–8^. Histones function by wrapping DNA into complex structures and, in doing so, control access to the genome. Histone proteins are modified by a wide-range of post-translational modifications (PTMs) that are dynamically regulated to control gene expression^9–11^. Histone mimicry allows viruses to disrupt the host cell’s ability to regulate gene expression and respond to infection effectively. Thus far, only a few cases of such mimicry have been observed and validated,^4,7^ and no known cases of histone mimicry have been reported within coronaviruses. Furthermore, there are few studies examining epigenetic changes associated with coronavirus infection^12–14^ and none yet published on SARS-CoV-2. However, recent work has demonstrated that SARS-CoV-2 only weakly induces type I and type III interferon expression indicating that it suppresses the innate antiviral host cell response^15–17^. While SARS-CoV-2 likely employs numerous mechanisms to dampen this response, we examined whether SARS-CoV-2 employs histone mimicry to disrupt histone regulation to better understand how it evades host cell antiviral responses.

To investigate whether histone mimicry is utilized by SARS-CoV-2, we first performed a bioinformatic comparison of all SARS-CoV-2 viral proteins^18^ with all human histone proteins (**Fig. S1a-b**). Most SARS-CoV-2 proteins are highly similar to those in the coronavirus strain that caused the previous major SARS-CoV outbreak with the notable exception of the proteins encoded by ORF3b and ORF8^19,20^. Remarkably, we detected an identical match between a region of the protein encoded by ORF8 (henceforth called Orf8) and critical regions within the histone H3 amino terminal tail (**Fig. 1a**). Furthermore, Orf8 aligns to a longer stretch of amino acids (6 identical, sequential amino acids) than any previously described and validated case of histone mimicry^4–7,21^ (**Fig. S1c**). Based on a crystal structure of Orf8, this region of the protein falls in a disordered region that lies on the surface of protein in an Orf8 monomer^22^. Most compellingly, the motif contains the ‘ARKS’ sequence, which is found at two distinct sites in the histone H3 tail (**Fig. 1a**) and is well-established as one of the most critical regulatory regions within H3. Both ARKS sites are modified with multiple PTMs, including methylation and acetylation at H3 Lysine 9 (H3K9me3 and H3K9ac) and at H3 Lysine 27 (H3K27me3 and H3K27ac). Strikingly, this amino acid stretch is absent from the previous SARS-CoV virus Orf8-encoded protein (both before and after a deletion generated two distinct peptides, Orf8a and Orf8b^23^) but present in Bat SARS-CoV-2 (**Fig. S1d**). This proposed histone mimicry motif is also a considerably closer match than a previously proposed histone mimic in protein E of SARS-CoV-2 (**Fig. S1c, S1e**)^24^. Finally, evidence from patients indicates that Orf8 is highly expressed during infection^25,26^ and transcriptional and proteomics analyses^27^ indicate that within 24 hours of infection, *ORF8* transcript is expressed at higher levels than histone H3 transcripts and Orf8 protein is expressed at over 20% of the level of the most abundant histone H3 protein (**Fig. S1f-g**). These findings are consistent with a model in which Orf8 acts as a histone mimic to disrupt regulation of ARKS sites on histone H3, providing a novel mechanism through which this highly divergent protein^28–30^ functions during infection.

**Figure 1.**
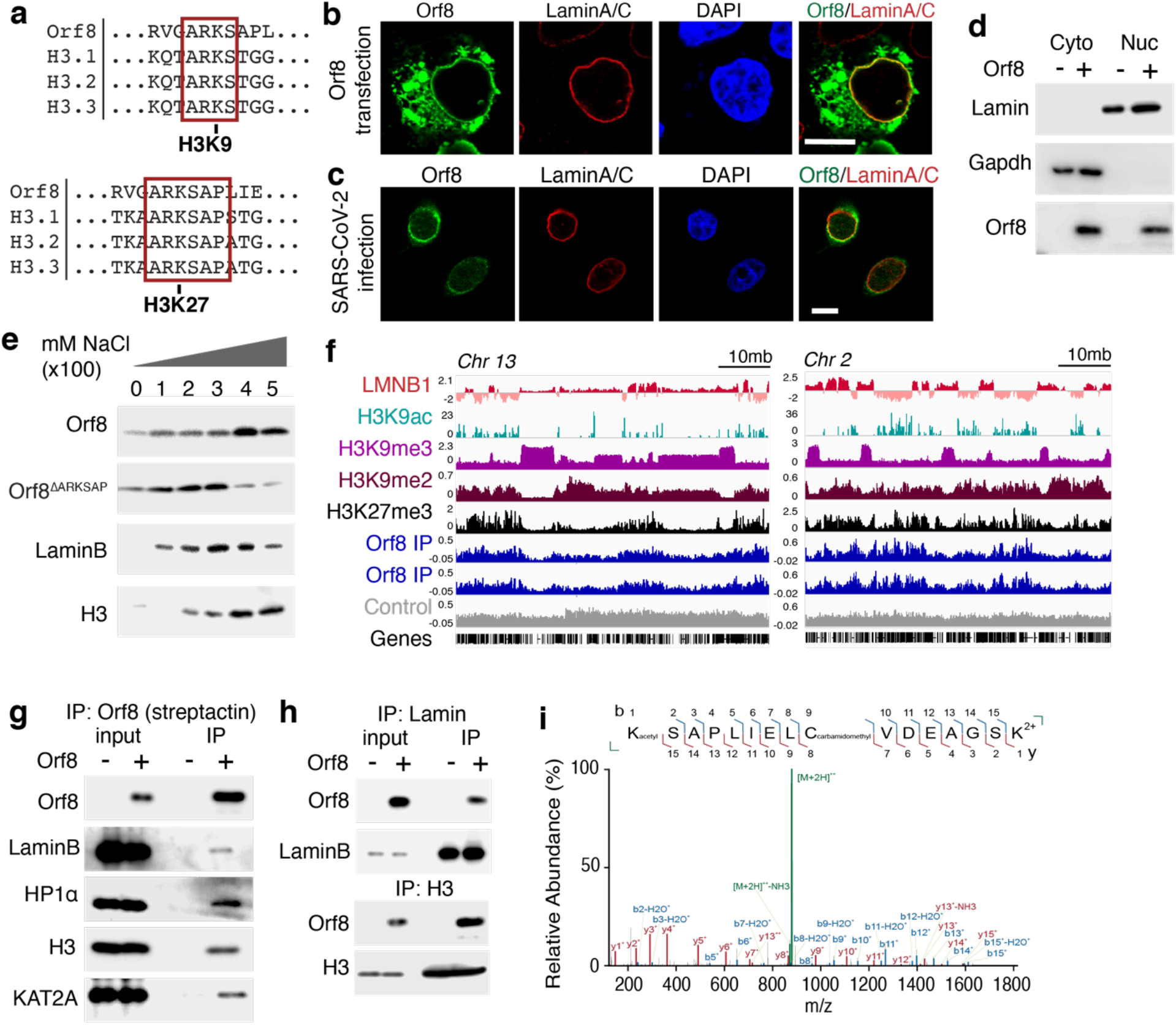
Orf8 associates with chromatin. (**a**) Orf8 contains an ARKS motif that matches the histone H3 tail regions surrounding the critical sites H3K9 and H3K27. (**b**) Staining of HEK cells transfected with Strep-Orf8 shows Orf8 is expressed in the cytoplasm and at the nuclear periphery colocalized with LaminA/C. (**c**) Orf8 and LaminA/C staining of SARS-CoV-2 infected A549^ACE^ cells at MOI=1, 48 hours post infection shows Orf8 is expressed at the nuclear lamina. (**d**). Subcellular fractionation of HEK cells transfected with Strep-Orf8 indicates Orf8 is present in the cytoplasm and nucleus. (**e**) Sequential salt extraction of HEK cells expressing a GFP control plasmid, Orf8, and Orf8^ΔARKSAP^. Orf8 is present in chromatin fractions containing LaminB and histone protein H3. (**f**) Gene tracks for Orf8 ChIP-sequencing normalized to input controls indicates similar binding patterns to H3K9me2 and H3K27me3. (**g**) Orf8 co-immunoprecipitates with Lamin complex-associated proteins including LaminB, HP1α, and H3, and KAT2A. ‘-’ indicates cells that are not expressing Orf8 for negative control IPs performed in parallel. (**h**) Reciprocal co-immunoprecipitation for Lamin and H3 confirm Orf8 binding. (**i**) Targeted mass spectrometry analysis of trypsin-digested Orf8 shows Orf8 is acetylated at lysine 52, at the site of the proposed histone mimic in Orf8. The intact 2+ charged peptide or precursor at 879.9508++ m/z was isolated and fragmented resulting in the MS/MS spectra shown. After fragmentation, the MS/MS spectra show unfragmented precursor (green) with matching product ions (b ions in blue, y ions in red) within 10ppm mass error. Each fragment’s intensity is given relative to the highest ion in the MS/MS spectra across the m/z range. The color, letter, and number of each fragment indicates the sequence that fragment contains within the larger peptide (top). Y fragments (red) indicate C-terminus matched fragments. B fragments (blue) indicate N terminus matched fragments. Scale bars = 10μM.

To determine whether Orf8 functions as a histone mimic, we began by examining its intracellular localization. While Orf8 does not have a well-defined nuclear localization sequence (NLS), it is 15kD in size and thus small enough to diffuse into the nucleus. We transfected HEK cells with Strep-tagged Orf8 and using immunofluorescence observed Orf8 in the cytoplasm and located at the periphery of the nucleus (**Fig. 1b**). This distribution matches a previous report^31^, although this study focused on a cytoplasmic role of Orf8. Given the observed expression pattern of Orf8, we next asked whether Orf8 is associated with Lamin. We found that Orf8 colocalized with LaminB1 and LaminA/C in cells transfected with Orf8 (**Fig. 1b, Fig. S2a-b**). Next, we infected an A549 lung cell line expressing the ACE receptor (A549^ACE^) with SARS-CoV-2, stained cells with an antiserum specific to Orf8 (**Fig. S2c-d**), and confirmed a similar expression pattern in infected cells (**Fig. 1c**). Finally, to confirm these findings through an independent approach we used cell fractionation and detected Orf8 in both the cytoplasm and the nucleus (**Fig. 1d**). Notably, while other functions have been proposed for Orf8^31–33^, the mechanisms through which it acts and what role it may play within the nucleus of host-cells are not fully understood.

We next tested whether Orf8 is associated with chromatin, using increasing salt concentrations to examine chromatin binding. We found that Orf8 dissociates from the chromatin fraction at salt concentrations similar to those at which Lamin and histones dissociate (**Fig. 1e**). In contrast, Orf8 containing a deletion of the ARKSAP motif (Orf8^ΔARKSAP^) (**Fig. S2e**) dissociates at lower salt concentrations indicating that the putative histone mimic site affects the strength of Orf8 binding to chromatin. Given that Orf8 appears to associate with Lamin and dissociates from chromatin at high salt concentrations containing histone proteins, we performed chromatin immunoprecipitation with high-throughput DNA sequencing (ChIP-seq) for Orf8 itself to determine if and where Orf8 associates with genomic DNA. We discovered that Orf8 shows substantial enrichment over input indicating that it is associated with chromatin (**Fig 1f**). Further comparisons with published datasets of histone modification ChIP-sequencing indicated that it is primarily found within genomic regions that are enriched in H3K27me3 and H3K9me2 but not H3K9me3 (**Fig. S3a**).

We next asked what specific proteins Orf8 associates with. Based on the localization of Orf8 to the periphery of the nucleus and its association with chromatin (observed using both biochemical and sequencing approaches), we began by examining Lamin-complex proteins. We found that Orf8 co-immunoprecipitated with LaminB1, histone H3, and HP1α, a protein associated with both Lamin and histones (**Fig. 1g**). Similarly, reciprocal co-immunoprecipitation for LaminB1 and Histone H3 confirmed Orf8 binding (**Fig. 1h**). Next, we tested whether Orf8 also co-immunoprecipitates with the histone-modifying enzymes that target the ARKS motif within Histone H3. We found that Orf8 is associated with the histone acetyl transferase KAT2A/GCN5 which targets H3K9 (**Fig. 1g**). Finally, we used mass spectrometry to identify additional binding partners beyond those found through a candidate approach (**Table S1)**. Hits included the HAT complex protein MORF4L, several zinc finger proteins, and the transcription factor SP2 which we confirmed by co-immunoprecipitation (**Fig. S3b**).

Because we observed that Orf8 can associate with KAT2A, we next used targeted mass spectrometry to determine whether the proposed Orf8 histone mimic site is modified similarly to histones. Using a bottom-up approach, Orf8 was purified from cells, reduced, alkylated, and digested. Separation with liquid chromatography was followed by parallel reaction monitoring mass spectrometry (LC-PRM-MS) targeting possible unmodified and modified forms of Orf8 commonly found on histones including serine phosphorylation, and lysine mono-methylation, di-methylation, tri-methylation, and acetylation. Of these targets, unmodified and acetylated lysine were identified. The acetylated peptide contained the +42 Da mass shift and demonstrated almost complete coverage of all possible product ions from the N-terminus containing the acetyl-lysine (b ions) as well as from the C-terminus (y ions). High resolution mass spectrometry differentiated the precursor from the trimethylated peptide and matched all product ions within 10 ppm mass error (**Fig. 1i, S3c**). This demonstrates that Orf8 is acetylated on the lysine within the proposed ARKS histone mimic site, similarly to histone H3. Notably, the presence of the acetylated lysine within the ARKSAP motif is likely incompatible with the dimerization of Orf8 which requires a covalent bond form at this residue^22^, and thus suggests that at least some portion Orf8 exists as a monomer within cells. In addition, this finding lends further support to Orf8 association with histone modifying enzymes such as KAT2A, suggesting that not only does Orf8 associate with proteins such as acetyltransferases, but is likely also modified by them similarly to histone H3.

Taken together, these findings demonstrate that Orf8 is well positioned to act as a histone mimic based on its localization at the nuclear lamina, its association with chromatin and chromatin-modifying enzymes, and its ability to retain the histone acetyltransferase KAT2A on chromatin.

To determine whether Orf8 does in fact act as a histone mimic, we sought to determine if it is capable of affecting chromatin regulation. We first examined whether Orf8 expression disrupts histone PTMs using an unbiased mass spectrometry approach. HEK cells were transfected with a control plasmid expressing GFP or with Orf8 containing a Strep tag. Transfected cells, identified by GFP fluorescence or with a Strep-Tactin conjugated fluorescent probe, were isolated using fluorescence-activated cell sorting (FACS) (**Fig. S4a**). Histones were purified through an acid-extraction method, and bottom-up unbiased mass spectrometry was performed to quantify all detected histone PTMs. Fitting with its potential role as a histone mimic, we found that numerous histone modifications were disrupted in response to Orf8 expression (**Table S2**). We focused on significantly disrupted histone PTMs with well-defined links to gene expression that contributed to at least 1% of the total peptide detected. Remarkably, we found numerous histone modifications associated with active gene expression were depleted, while histone modifications associated with chromatin compaction or transcriptional repression were increased in cells expressing Orf8 (**Fig. 2a**). In particular, modifications within the H3 ARKS motifs were highly disrupted. Peptides containing dual H3K9ac and H3K14ac, both well-established PTMs linked to active gene expression, were decreased by Orf8 expression. Conversely, the peptides containing H3K9 methyl modifications (H3K9me2 and H3K9me3), as well as peptides containing H3K27 methylation (both on the canonical H3 and variant H3.3 histone, H3K27me3 and H3.3K27me3), were robustly increased in response to Orf8. These data support a role for Orf8 as a putative histone mimic and demonstrate that it is capable of disrupting histone PTM regulation at numerous critical sites within histones.

**Figure 2.**
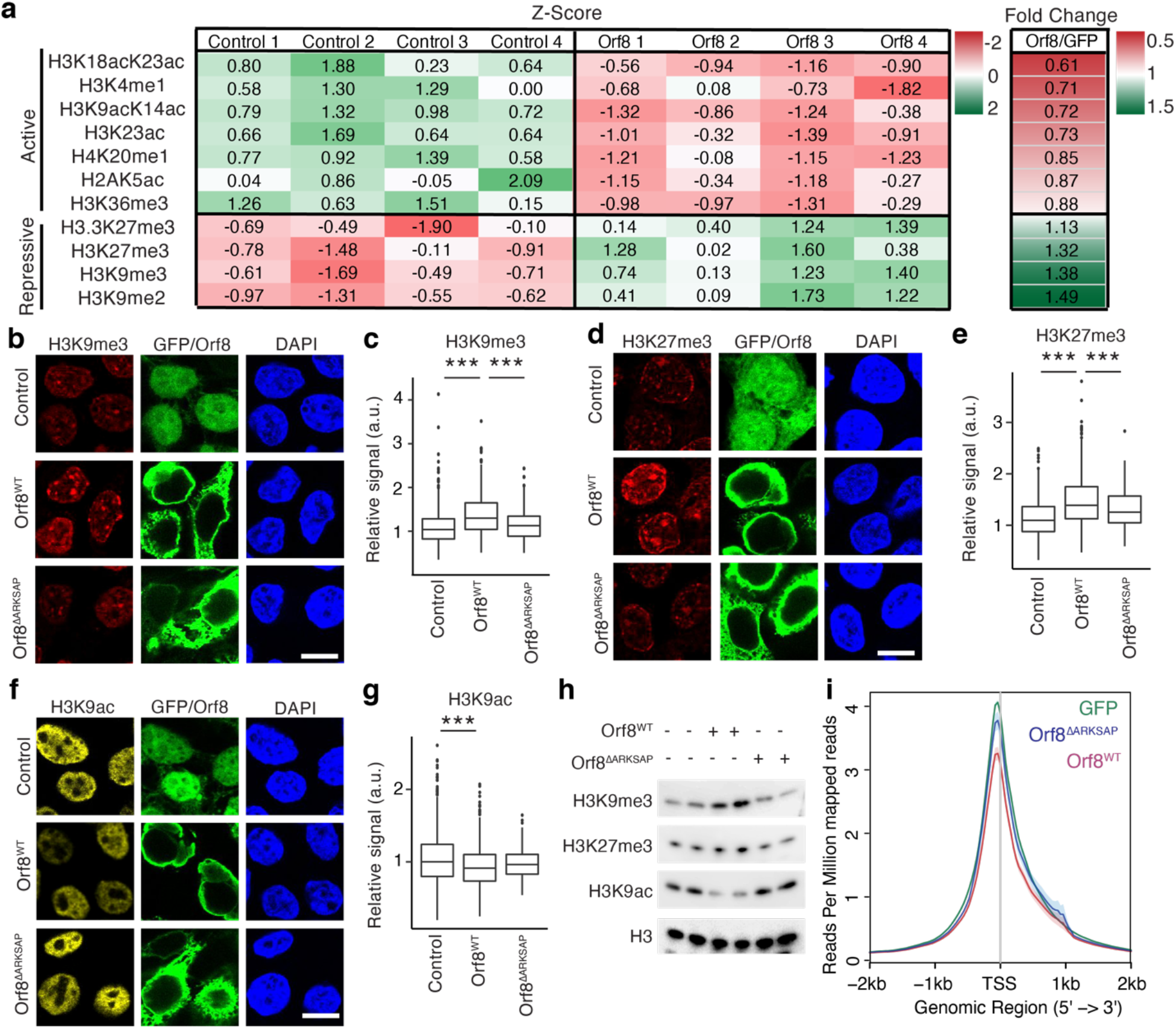
Orf8 function in hPTM regulation. (**a**) Mass spectrometry analysis of histone PTMs in control (GFP) or Orf8 expressing HEK293T cells isolated by FACS. Z-score and fold changes are shown for modifications that are significantly changed in response to Of8 expression, were detected over a minimal threshold of 1% of the total peptide abundance, and have well-established functions. Full results and raw data are shown in Table S1. (**b-g**) HEK293Tcells transfected with GFP or Strep-Orf8 show that Orf8 expression increases H3K9me3 (b-c) and H3K27me3 (d-e) while decreasing H3K9ac (f-g). Conversely, Orf8 with a deletion of the histone mimic site ARKSAP (Orf8^ΔARKSAP^) does not affect these histone PTMs. N = 614 (GFP), 497 (Orf8), 170 (Orf8^ΔARKSAP^) cells for H3K9me3; 616, 550, 154 cells for H3K27me3; 666, 568, 170 cells for H3K9ac compiled from 3 independent transfections. ***, p<0.001, 1-way ANOVA with post-hoc 2-sided t-test and Bonferroni correction. (**h**) Western blot analysis of histones isolated from FAC sorted transfected cells. (**i**) ATAC-seq of HEK293Tcells expressing GFP or Orf8 isolated by FACS. Reads per million mapped surrounding the transcription start site (TSS) of all expressed genes are averaged. N = 2 independent replicates. Scale bars = 10μM.

To confirm mass spectrometry findings through an independent approach, we again transfected HEK cells with Orf8 and examined global changes in modifications which lie within the proposed histone mimic motif and which mass spectrometry data indicated are disrupted by Orf8. We used immunofluorescent imaging to measure methylated and acetylated H3K9 and H3K27 as well as Orf8 protein to ensure that analyzed cells contained equivalent levels of various Orf8 constructs (**Fig. S4b**). We found that cells expressing Orf8 exhibited increased H3K9me3 and H3K27me3 and decreased H3K9ac staining compared to control plasmid transfected cells (**Fig. 2b-g**). Similar to mass spectrometry findings, we did not detect significant global changes in H3K27ac using this method (**Fig. S4c**), potentially due to low basal levels of H3K27ac. To determine whether these effects are due to the proposed histone mimic site within Orf8, we generated a deletion construct lacking the ARKSAP histone mimic site (Orf8^ΔARKSAP^). While Orf8^ΔARKSAP^ was expressed at similar levels to Orf8 (**Fig. S4b)**, it did not increase H3K9me3 or H3K27me3, and had a non-significant intermediate effect on H3K9ac (**Fig. 2b-g**). Next, we examined an acquired mutation in Orf8 commonly found in SARS-CoV-2 strains, S84L (Orf8^S84L^). This site is unlikely to affect the protein stability^34,35^ and lies outside the histone mimic region and thus is not expected to affect its ability to regulate histone PTMs. We found that Orf8^S84L^ also increased H3K9me3 and H3K27me3, while decreasing H3K9ac (**Fig. S4e-g**), indicating that, as predicted, this common mutation does not alter the histone mimic function of Orf8. Similarly, a 6 amino acid deletion in another unstructured region of Orf8 that with similar amino acid make-up but in a different sequence (AGSKSP) as the histone mimic does not affect the ability of Orf8 to disrupt histone regulation (**Fig. S4g**).

We next sought to confirm these findings using independent methods. To ensure equal levels of expression Orf8 and Orf8^ΔARKSAP^, we isolated transfected cells by FACS (**Fig. S5a**). We then isolated histones through acid extractions and examined histone modifications by western blot. We confirmed that Orf8 increased H3K9me3 and H3K27me3 and deceased H3K9ac in an ARKSAP-dependent manner (**Fig. 2h**). Next, we isolated transfected cells (**Fig. S5b**) and performed CUT&TAG sequencing of H3K9ac. We found that Orf8, but not Orf8^ΔARKSAP^, deceased H3K9ac (**Fig. S5c**). Based on the observed pattern of histone PTM disruptions, we hypothesized that Orf8 expression may also decrease chromatin accessibility. We isolated transfected cells using FACS (**Fig S5d**), and performed Assay for Transposase-Accessible Chromatin with high-throughput sequencing (ATAC-seq) to assess changes in open and closed chromatin. Orf8, but not Orf8^ΔARKSAP^, resulted in robust decreased chromatin accessibility (**Fig. 2i**). Further, both H3K9ac and chromatin accessibility changes occurred globally regardless and were particularly evident in mid to highly expressed genes (**S5e-h**). To determine how these global changes to chromatin affect gene expression, we again isolated transfected cells using FACS and used RNA-sequencing (RNA-seq) to define differentially expressed genes (**Fig. S6a**). We found that both Orf8 and Orf8^ΔARKSAP^ caused robust changes in gene expression and shared a subset of differentially expressed genes. However, the presence of the histone mimic motif resulted in less dynamic gene expression changes in cells in response to expression of exogenous Orf8 (**Fig. S6b-d**). In addition, distinct gene groups were also differentially expressed between these two forms of Orf8, with Orf8 causing decreases in gene expression compared to Orf8^ΔARKSAP^ (**Fig. S6e**), particularly of mid to highly expressed genes and genes associated with DNA regulation and cellular response pathways (**Fig. S6f-i, Table S3**) These data define a role for Orf8 in disruption of host cell histone PTMs through a novel case of histone mimicry of the ARKS motifs in H3.

Having shown that Orf8 alone is sufficient to disrupt chromatin regulation, we next examined the effect of Orf8 on histone PTM regulation in the context of viral infection. In order to test whether effects on PTM regulation are due to Orf8, we prepared a recombinant mutant SARS-CoV-2 with a deletion of Orf8 (SARS-CoV-2^ΔOrf8^) using the cDNA reverse genetics system.^36,37^ We infected A549^ACE^ cells with SARS-CoV-2 or SARS-CoV-2^ΔOrf8^ at an MOI of 1 and compared the level of viral genomes and infectious virus production in the presence and absence of Orf8. Due to the overexpression of the ACE receptor, these cells are readily and rapidly infected by SARS-CoV-2 and thus provide an ideal system in which to compare the early cellular response to different forms of a virus without the complication of different rates of infection. We found similar levels of viral genome copies in SARS-CoV-2 and SARS-CoV-2^ΔOrf8^ infected cells at 24 and 48 hours after infection (**Fig. S7a**). Additionally, no differences in viral titer were observed at 24 hours and only subtle differences were detected at 48 hours after infection through plaque assays (**Fig. 3a**). Critically, this allows for a direct comparison of these two viruses at early time points after infection in A549^ACE^ cells without the confounding effects of differences in the amount of virus present. We therefore infected A549^ACE^ cells with SARS-CoV-2 or SARS-CoV-2^ΔOrf8^ or performed a mock infection and used ChIP-sequencing with ChIP-RX normalization to allow for the detection of global as well as site specific changes in histone PTMs. Remarkably, we found that SARS-CoV-2 infection results in robust increases in H3K9me3 and H3K27me3 (**Fig. 3b-c**), mirroring the effects of Orf8 expression. However, SARS-CoV-2^ΔOrf8^ infection more closely resembled mock infected cells than wildtype SARS-CoV-2 infected cells indicating that the effect of SARS-CoV-2 on repressive histone modifications is largely due to Orf8 expression. Conversely, SARS-CoV-2 infection resulted in a global decrease in H3K9ac (**Fig. 3d**), again matching the effects of Orf8 expression. Orf8 deletion not only reduced the effect of SARS-CoV-2 on H3K9ac, but SARS-CoV-2^ΔOrf8^ infection resulted in an increase in H3K9ac compared to mock infection. Finally, we used ATAC-seq to examine chromatin compaction and found that wildtype SARS-CoV-2 decreased chromatin accessibility and this effect was largely lost with deletion of Orf8 (**Fig. 3e**). These effects were evident at both the global level as well as at specific genes involved in viral response pathways (**Fig. 3f, S7b**).

**Figure 3.**
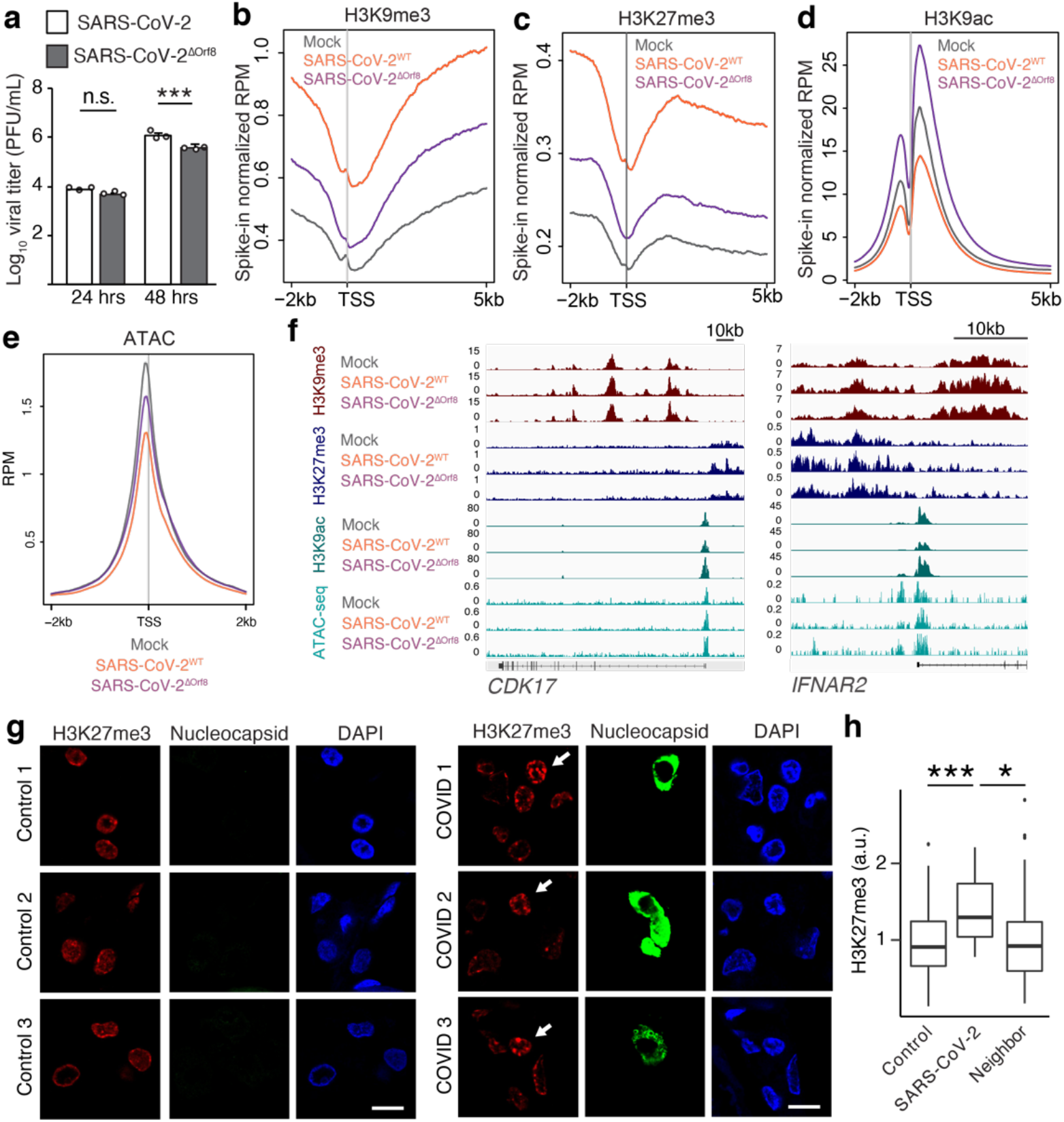
SARS-CoV-2 infection affects histone PTMs. (**a**) qRT-PCR analysis of expression of viral titer in A549^ACE^ pulmonary cells at 24 (a) and 48 (b) hours after infection with SARS-CoV-2^WT^ or SARS-CoV-2^ΔOrf8^ at MOI=1. n = 3 replicates from infection done in parallel. (**b-d**) ChIP-RX for H3K9me3 (b), H3K27me3 (d) and H3K9ac (d) of A549^ACE^ cells with SARS-CoV-2^WT^, SARS-CoV-2^ΔOrf8^, or mock infection at MOI=1, 48 hours after infection. n = 3. (**e**) ATAC-seq of A549^ACE^ cells with SARS-CoV-2^WT^, SARS-CoV-2^ΔOrf8^, or mock infection at MOI=1, 48 hours after infection, n = 2 for ATAC. (**f**) ChIP and ATAC-seq gene tracks of genes in signaling pathways and cytokines relevant to viral response. (**g**) Postmortem COVID-19 patient lung tissue stained for H3K9me3 and Nucleocapsid protein to identify SARS-CoV-2 infected cells. (**h**) Quantification of H3K9me3 in infected cells compared to neighboring cells from the same tissue slice. N = 12 SARS-CoV-2 infected cells and 131 uninfected neighboring cells from 3 COVID patient samples and 60 cells from 3 control patients. *,p<0.05, ***, p<0.001, 1-way ANOVA with post-hoc 2-sided t-test and Bonferroni correction. Scale bars = 10μM.

While our data indicated that the differences in SARS-CoV-2 and SARS-CoV-2^ΔOrf8^ are unlikely to be due to any difference in viral replication (**Fig. 3a, S7a**), we sought to further confirm these finding using an approach that is independent of the number of cells infected. We used immunohistochemistry to stain for histone modifications of interest and used dsRNA staining to identify and specifically examine infected cells. We found that at 24 hours after infection, cells infected with SARS-CoV-2 had increased H3K9me3 and H3K27me3 and decreased H3K9ac compared to either mock infected cells or uninfected neighboring cells (**Fig. S7c-h**). As observed in ChIP-sequencing data, this effect was largely lost with deletion of Orf8.

Lastly, to determine whether similar effects also occur in the context of a patient population, we obtained postmortem lung tissue samples from three COVID-19 patients and matched controls. We stained tissue for H3K9me3 and for SARS-CoV-2 nucleocapsid protein to identify infected cells. We found that in all patient samples, infected cells showed increased H3K9me3 staining compared to neighboring cells within the same tissue, as well as compared to control tissue (**Fig. 3g-h, S7i**). While sample availability limits the conclusions that can be drawn from this assay, this finding indicates that histone PTMs are also disrupted in patients with severe COVID-19 disease. In summary, we found that the effects of SARS-CoV-2 infection on histone PTMs and chromatin compaction require Orf8 expression and mirror the ARKSAP-dependent effects of Orf8.

Next, we examined how Orf8 affects the transcriptional response to SARS-CoV-2 infection. We used RNA-seq to examine A549^ACE^ cells at 24-hours after infection to ensure equivalent viral particles and genome copies of SARS-CoV-2 and SARS-CoV-2^ΔOrf8^ (**Fig. 3a, S7a**). As further confirmation of equivalent viral infectivity at this time point, we detected no significant differences in the number of reads that map to the SARS-CoV-2 versus human genomes between SARS-CoV-2 and SARS-CoV-2^ΔOrf8^ (**Fig. S8a**). As expected, the only difference in expression of individual SARS-CoV-2 genes between the viruses was for Orf8 (**Fig. S8b**). However, within the wildtype virus, Orf8 transcript was highly expressed and found at greater levels than many of the transcripts that encode the human proteins that it interacts with (**Fig. S8c**). Interestingly, SARS-CoV-2^ΔOrf8^ resulted in more differentially expressed genes than wildtype SARS-CoV-2 (**Fig. 4a-b**) indicating that Orf8 partly represses the dynamic transcriptional response to infection. Direct comparison between the viruses (**Fig. 4c, S8d-e**) and GO analysis (**Fig. S9, Table S4**) indicated that different groups of genes were disrupted by the two viruses, with SARS-CoV-2^ΔOrf8^ infection resulting in an induction of numerous genes relevant to the host cell response to infection such as interferon response genes, cytokine-encoding genes, and other critical signaling pathway genes. By 48 hours after infection, both viruses occupied the vast majority of the mapped reads (**Fig. S10a**) and both resulted in robust changes in gene expression compared to mock infected cells. At this time point, the wildtype virus surpasses the SARS-CoV-2^ΔOrf8^ transcriptional response (**Fig. S10b-d**) potentially due to the slightly greater number of infectious particles generated by wildtype SARS-CoV-2 by this time point (**Fig. 3a**). Finally, we compared the changes in histone PTMs detected through ChIP-sequencing to differentially expressed genes. We found that changes in H3K9ac most closely tracked with gene expression changes in response to infection with repressive marks showing similar global changes in all gene groups (**Fig. S11**). Notably, these data further support recent findings indicating SARS-CoV-2 results in a limited transcriptional response^15,16,38^ and demonstrate that Orf8 is at least partly responsible for repressing the transcriptional response at early time points after infection. This fits with its role in decreasing activating histone modifications and promoting repressive modifications and chromatin compaction.

**Figure 4.**
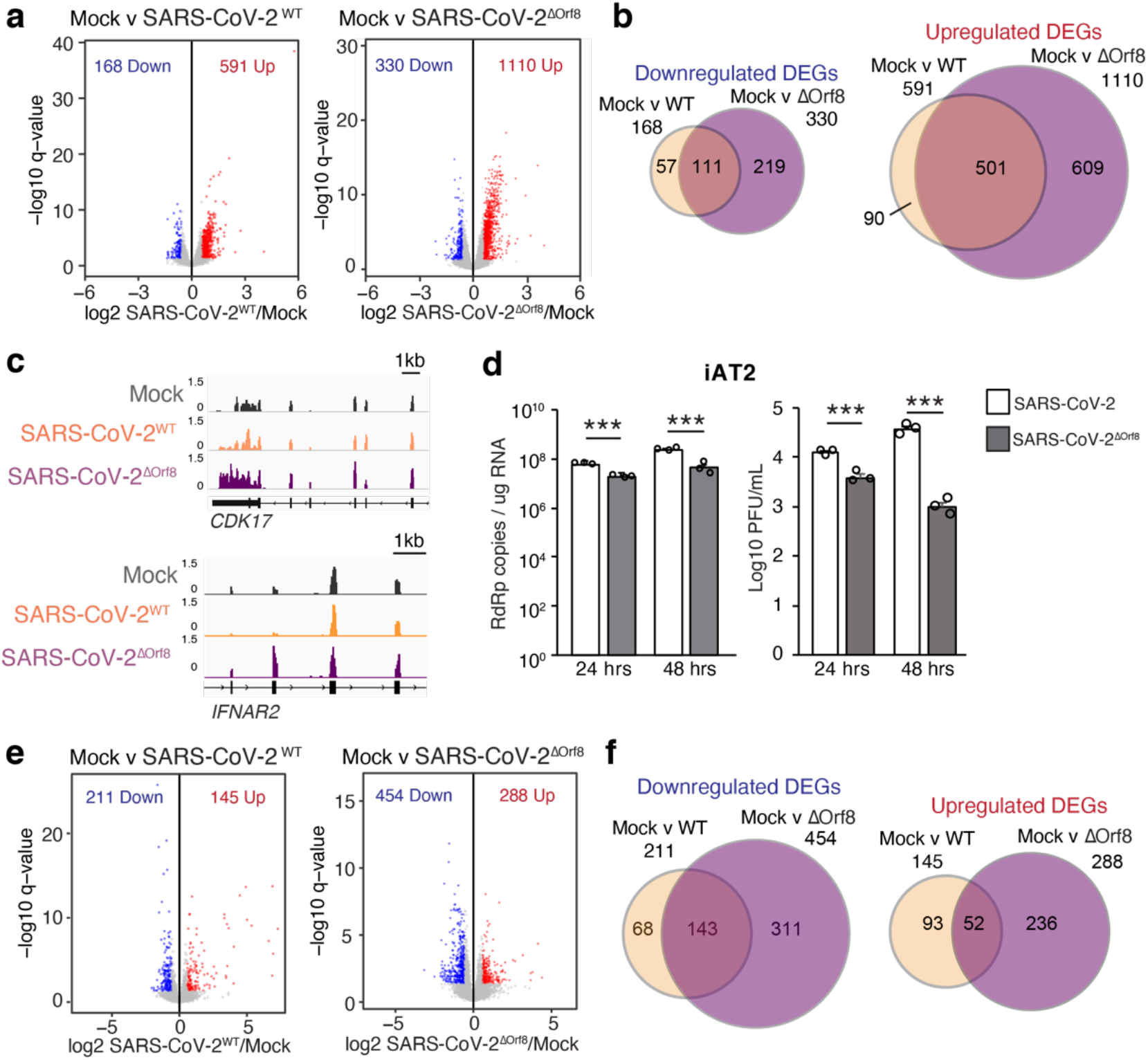
Orf8 affects gene expression and viral replication during SARS-CoV-2 infection. (**a**) Differential gene expression analysis by RNA-seq of A549^ACE^ cells 24 hours after SARS-CoV-2^WT^, SARS-CoV-2^ΔOrf8^, or mock infection at MOI=1. Significantly differentially expressed genes (relative to mock infection) are shown in blue (down) and red (up). N = 3. (**b**) Overlap of differentially expressed genes in response to SARS-CoV-2^WT^ and SARS-CoV-2^ΔOrf8^ infection. (**c**) Gene tracks of examples of genes that show chromatin disruptions in response to SARS-CoV-2^WT^ but not SARS-CoV-2^ΔOrf8^ infection. (**d**) qRT-PCR analysis of expression of SARS-CoV-2 gene RdRp and plaque assay analysis of viral titer in induced pluripotent stem cell-derived lung alveolar type 2 (iAT2) pulmonary cells at 24 or 48 hours after infection with SARS-CoV-2^WT^ or SARS-CoV-2^ΔOrf8^ at MOI=1. n = 3 replicates from infection done in parallel. (**e**) Differential gene expression analysis by RNA-seq of iAT2 cells 48 hours after SARS-CoV-2^WT^, SARS-CoV-2^ΔOrf8^, or mock infection at MOI=1. Significantly differentially expressed genes (relative to mock infection) are shown in blue (down) and red (up). N = 3. (**f**) Overlap of differentially expressed genes in response to SARS-CoV-2^WT^ and SARS-CoV-2^ΔOrf8^ infection. ***, p<0.001, 2-way ANOVA with post-hoc 2-sided t-test and Bonferroni correction.

Given the robust effects of Orf8 deletion on host-cell chromatin regulation and the transcriptional response to infection, we sought to test whether Orf8 mediates the replication of SARS-CoV-2 using a cell type more physiologically relevant to human infection than A549 cells engineered to express high levels of the ACE receptor. To this end, we infected induced human pluripotent stem cell-derived lung alveolar type 2 (iAT2) pulmonary cells with SARS-CoV-2 and SARS-CoV-2^ΔOrf8^ (with an MOI of 1). Strikingly, we observed differences in both copies of viral genes and the proportion of RNA-seq reads mapped to the viral genome at both 24 and 48 hours after infection (**Fig. 4d, S12a**). Even more striking, viral titers measured through plaque assays demonstrate that SARS-CoV-2^ΔOrf8^ generated fewer infectious particles than wildtype SARS-CoV-2 even by 24 hours with robust differences detected at 48 hours (**Fig. 4d**). We further confirmed these findings with independently derived and infected iAT2 cells (**Fig. S12b**). These results demonstrate that Orf8 allows for increased replication in human lung epithelial cells.

Finally, we used RNA-seq to examine the differences in the transcriptional response of iAT2s to SARS-CoV-2 or SARS-CoV-2^ΔOrf8^ infection. We detected very few gene expression changes at 24 hours after infection (**Fig. S12c-e**), fitting with a slower infection rate in iAT2 cells compared to A549^ACE^ cells. We next examined gene expression changes at 48 hours after infection at which point the number of viral transcripts more closely aligns to that seen in A549^ACE^ cells at 24 hours (**Fig. S12a, S8a**). We found that similar to A549^ACE^ cells, SARS-CoV-2^ΔOrf8^ infection resulted in greater numbers of gene expression changes than wildtype SARS-CoV-2 (**Fig. 4e-f, S12f**). This difference is even more notable given that it occurs despite the decreased viral genome copies present and decreased viral particles produced by SARS-CoV-2^ΔOrf8^ infection. The effects of deletion of accessory proteins from SARS-CoV-2 in a transgenic mouse model appear complex, with Orf8 loss causing decreases in replication and viral load and limited effects on survival.^39^ However, new data from COVID-19 human patients examined a rare 382-nucleotide deletion variation in SARS-CoV-2 found in Singapore that results in the loss of a small portion of ORF7B and the majority of the ORF8 gene. This work found that this SARS-CoV-2 variant is associated with a milder infection in COVID-19 patients and an improved interferon response^40,41^. Our findings in a human iPSC line point toward the loss of Orf8 as a possible cause of these differences and provide a mechanism underlying the role of Orf8 in promoting SARS-CoV-2 virulence within the patient population.

The work described here identifies a novel case of histone mimicry during infection by SARS-CoV-2 and defines a mechanism through which SARS-CoV-2 acts to disrupt host cell chromatin regulation. We found that the protein encoded by the SARS-CoV-2 ORF8 gene contains an ARKS motif and that Orf8 expression disrupts histone PTM regulation. Orf8 is associated with chromatin-associated proteins, histones, and the nuclear lamina and Orf8 is itself acetylated within the histone mimic motif similarly to histones. Further, our data indicate that both Orf8 expression and SARS-CoV-2 infection result in global changes in histone regulation in multiple cell lines as well as patient tissue. Critically, our data indicate that Orf8 is required for the ability of SARS-CoV-2 to disrupt host cell chromatin and is sufficient to mediate these effects when exogenously expressed. In addition, these effects are reliant on the ARKSAP histone mimic motif within Orf8 protein. Notably, this work provides a molecular basis for the 2020 discovery that patients infected with a form of SARS-CoV-2 containing a deletion in the gene encoding Orf8 mount a stronger adaptive immune response^41^ and improved outcomes^40^.

Together, these findings provide a novel function of the highly divergent SARS-CoV-2 protein Orf8 and define the epigenetic disruptions that occur in response to infection. Notably, the role of Orf8 in chromatin disruption early in infection is not inconsistent with other proposed roles of Orf8 in other cellular compartments or at later stages of infection^31,33,35,42,43^. However, this work presents the first link between a specific SARS-CoV-2 encoded protein and the epigenetic disruptions that occur in response to infection and provides a mechanistic explanation for mounting evidence^4,40,44,45^ that epigenetic disruptions contribute to the severity of COVID-19.

Given that many epigenetic pathways and histone modifying enzymes are druggable, in many cases with therapeutics already approved for use in humans, this work suggests potential avenues for the development of treatments that target epigenetic pathways. Finally, these data have critical implications for our understanding of emerging viral strains carrying deletions and mutations in the ORF8 gene and COVID-19 pathogenesis in patients.

## Author contributions

J. Kee designed, performed, and analyzed the majority of the experiments. S. Thudium generated cells, samples, and DNA constructs. K. Palozola performed ATAC-seq and generated samples for histone PTM analysis. K. Gladstad and Z. Zhang performed and analyzed ChIP-sequencing experiments. J. Cesare performed and analyzed mass spectrometry experiments with guidance from B.A. Garcia. X. Zhang, X. Xie, and P.Y. Shi generated the Orf8 deletion virus. Y. Lan provided bioinformatic analysis. F.L. Cardenas and R. Truitt generated iAT2 cells with guidance from W. Yang and E. Morrisey. D.N. Kotton and K.D. Alysandratos provided stem cell lines. D. Renner and Y Li performed the SARS-CoV-2 viral infections and analysis of viral infections. S.L. Berger provided input and expertise and lead ChIP-sequencing studies. S.R. Weiss provided input and expertise and lead viral work. E. Korb lead the project and wrote the manuscript.

## Acknowledgments

We thank and acknowledge M. Weitzman for feedback and suggestions, A. Stout for microscopy support, C. Comar for viral infections, R. Jain and P. Shaw for protocols, S. Wolf for supervision, and M. Feldman and K. Montone for providing patient samples. J.K. was supported by the Penn Center for Coronavirus and Other Emerging Pathogens. E.K was supported by the Sloan Research Fellowship, the Klingenstein-Simons Fellowship, the NARSAD Young Investigator Award, and NIH grants R00MH111836 and 1DP2MH129985. Viral work was supported by NIH grant RO1-AI140442 supplement for SARS-CoV-2 and funds from Penn Center for Coronavirus Research and Other Emerging Pathogens. D. R. was supported in part by T32 AI055400. R.T. and W.Y. were supported in part by institutional funds from the University of Pennsylvania Perelman School of Medicine to the iPSC Core. iPSC cell line generation and sharing were supported by NIH grants NO1 75N92020C00005 and U01TR001810. Mass spectrometry was supported by AI118891. P.-Y.S. was supported by NIH grants AI134907 and UL1TR001439, and awards from the Sealy Smith Foundation, the Kleberg Foundation, the John S. Dunn Foundation, the Amon G. Carter Foundation, the Gillson Longenbaugh Foundation, and the Summerfield Robert Foundation.

## Conflict of Interest

The P.Y. Shi laboratory has received funding support in sponsored research agreements from Pfizer, Gilead, Novartis, Merck, GSK, IGM Biosciences, and Atea Pharmaceuticals. P.Y.S. is a member of the Scientific Advisory Boards of AbImmune and is Founder of FlaviTech.

Susan R Weiss is on the Scientific Advisory Board of Ocugen, Inc and immunome, Inc.

**Supplemental Figure 1.**
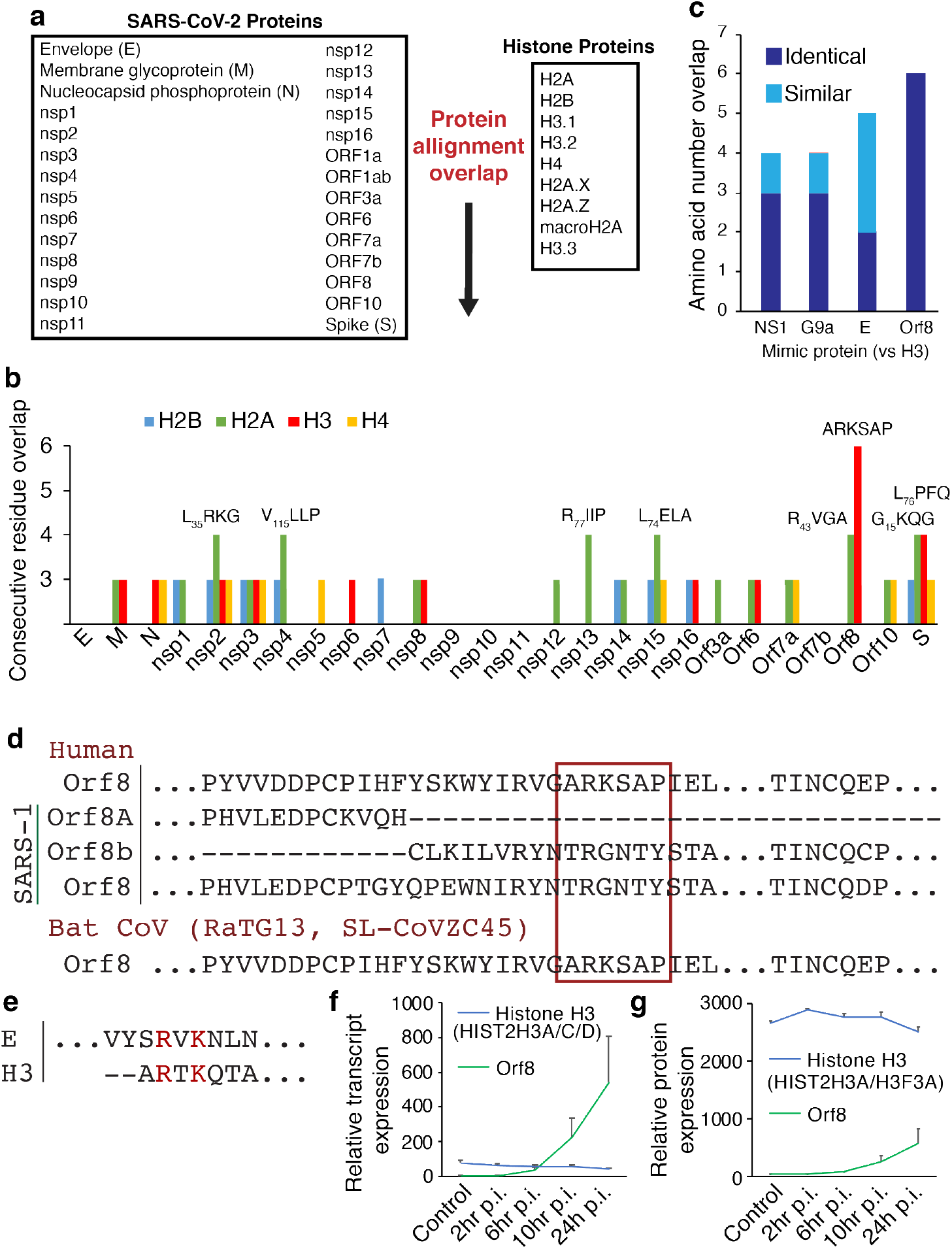
Orf8 histone mimicry and localization. (**a**) Alignments performed to identify putative histone mimic sites within the SARS-CoV-2 genome. (**b**) The number of exact sequential overlapping amino acids found between SARS-CoV-2 proteins and histone proteins. (**c**) The overlap of Orf8 and H3 compared to other proposed cases of histone mimicry. Exact amino acid overlap shown in dark blue and structurally similar amino acid overlap shown in light blue. NS1 is from Influenza A H3N2. Protein E is a previously proposed mimic in SARS-CoV-2. G9a is a human protein that mimics H3. (**d**) The ARKS motif is present in Bat SARS-CoV Orf8 but is not found with SARS-CoV Orb8a/b or the SARS-CoV precursor Orf8 before a mutation generated two distinct proteins. (**e**) Previously proposed histone H3 mimic in SARS-CoV-2 protein E. (**f-g**) Orf8 transcript (f) and protein (g) expression in SARS-CoV-2 infected Caco-2 cells from published datasets. MOI=1.

**Supplemental Figure 2.**
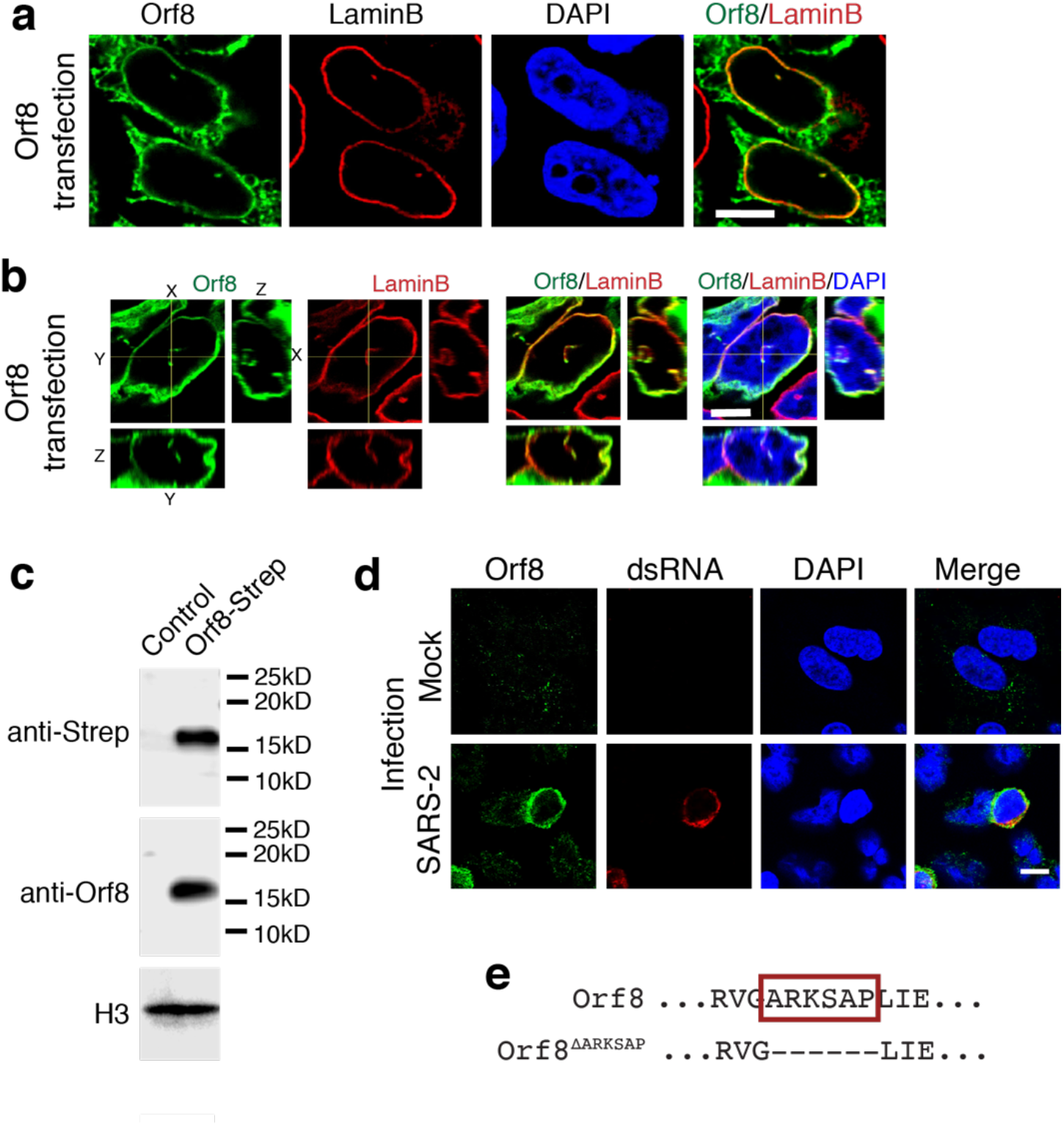
Orf8 localization. (**a**) HEK cells expressing Orf8 co-stained with Lamin B and streptactin to detect Orf8. (**b**) Rotation of z-stacks (right and bottom panel for each stain) indicate colocalization of Orf8 and LaminB throughout the nucleus. (**c**) Orf8 antiserum specifically detects Strep-tagged Orf8 by western blot In HEK cells transfected with Strep-Orf8. (**d**) Orf8 antiserum specifically stains infected A549^ACE^ cells 48 hours after SARS-CoV-2 infection with no staining observed in mock infection. (**e**) Deleted construct used to test effects of ARKSAP motif in Orf8. Scale bar = 5μM in a-b, 10μM in d.

**Supplemental Figure 3.**
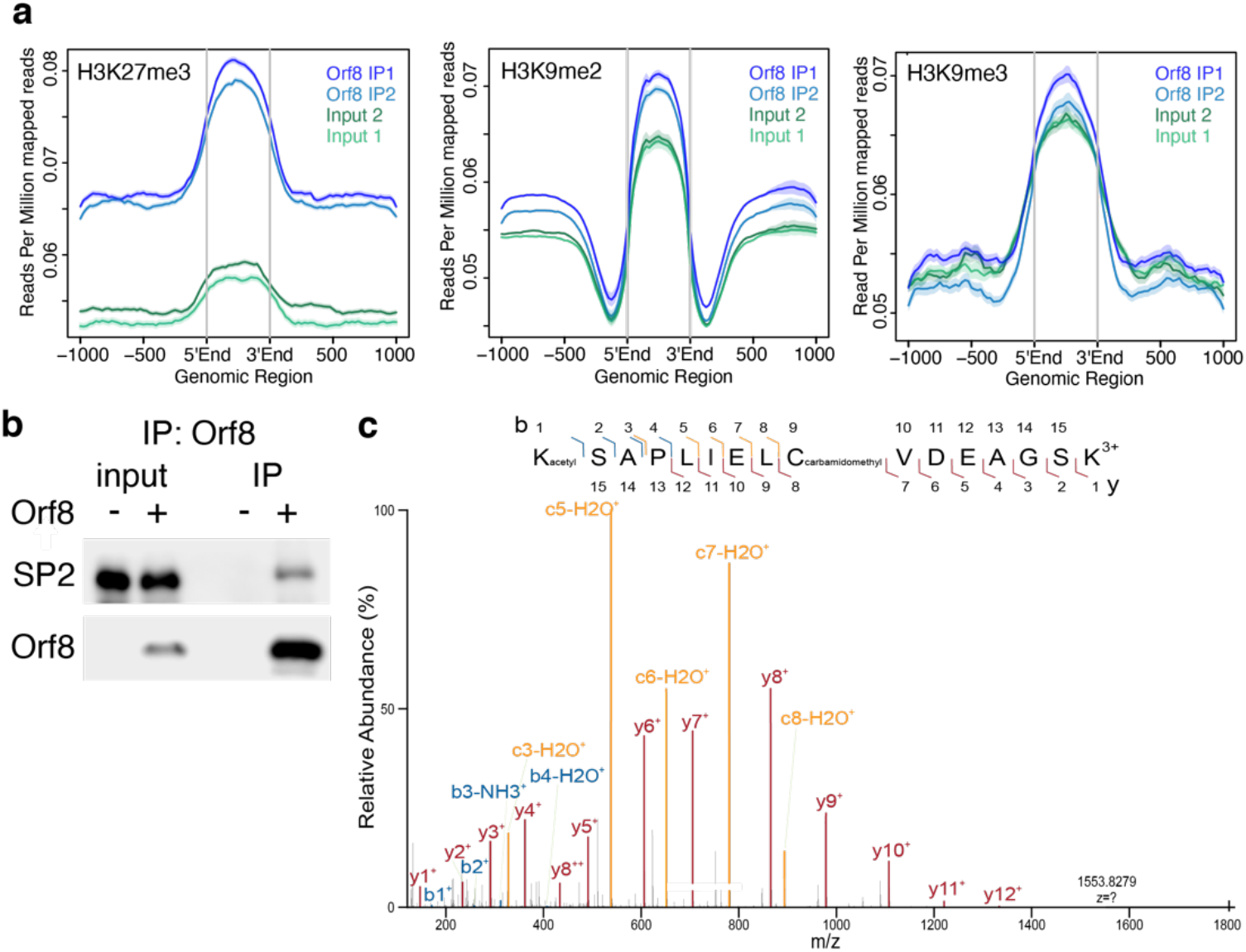
(**a**) Orf8 is enriched in genomic regions with high H3K27me3 and H3K9me2 relative to input controls with lower enrichment at regions with high H3K9me3. (**b**) Confirmation of mass spectrometry analysis of binding partner, transcription factor SP2. Orf8 was immunoprecipitated with Streptactin beads and resulting lysates were probed for SP2 and Streptactin. (**c**) Targeted mass spectrometry analysis of trypsin-digested Orf8 of the peptide containing the proposed histone mimic in Orf8. Orf8 acetylation at K52 is detected in the 3+ charged peptide by targeted mass spectrometry. MS/MS spectra with matching product ions (b ions in blue, y ions in red, c ions in yellow) within 10ppm mass error. Scale bar = 10μM.

**Supplemental Figure 4.**
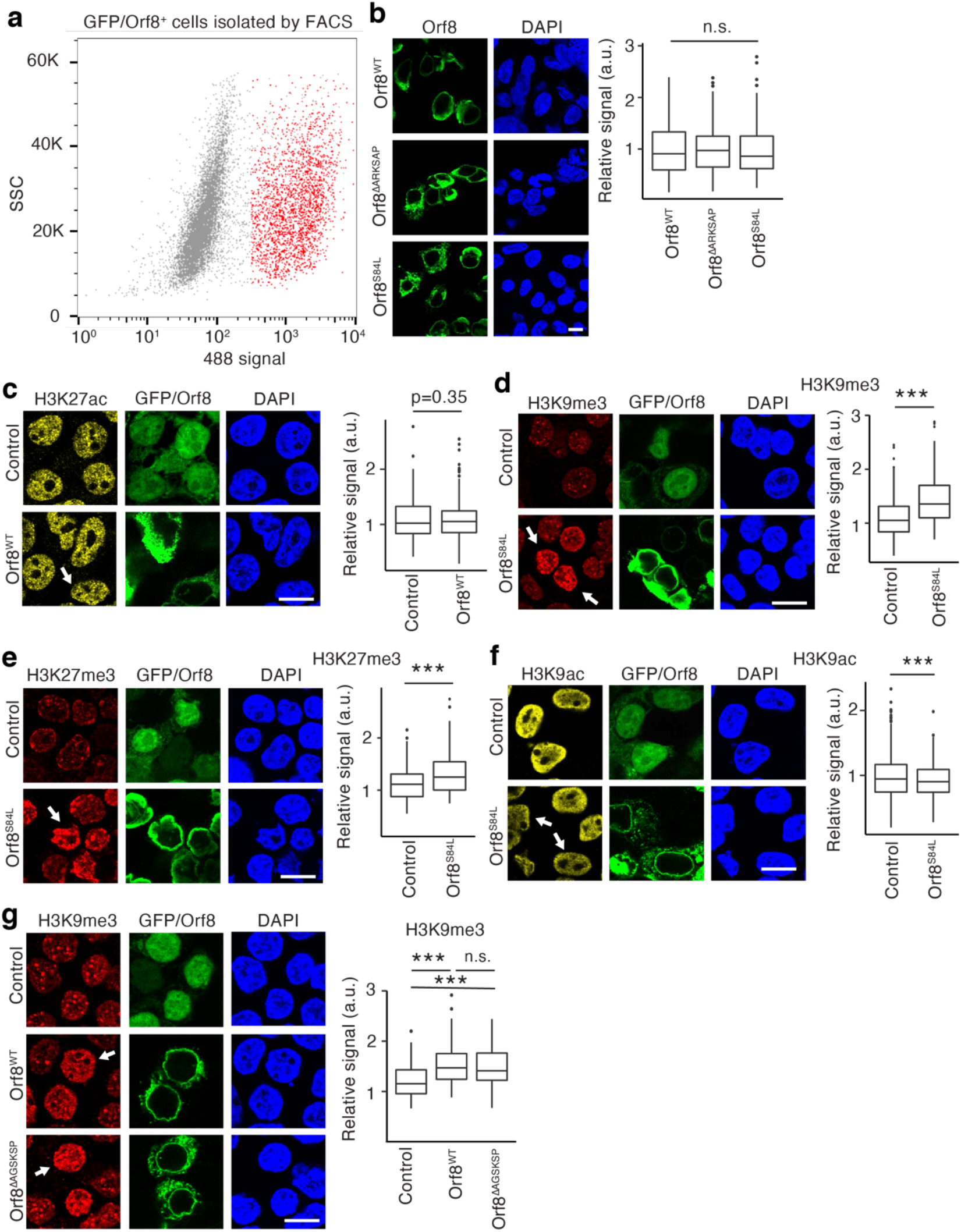
Orf8 and Orf8-S84L effects on histone PTMs. (**a**) Example of FACS used to isolate transfected cells expressing GFP, Orf8, or Orf8^ΔARKSAP^ and ensure equivalent levels of Orf8 and Orf8^ΔARKSAP^. Shown is an example for Orf8 with red indicating cells isolated for analysis. (**b**) Orf8 constructs tested show similar levels of expression in HEK cells. N = 137 (Orf8), 87 (Orf8^ΔARKSAP^), 120 (S84L) from 4 independent transfections. (**c**) Orf8 does not significantly affect H3K27ac. N = 616 (GFP), 550 (Orf8) from 3 independent transfections. (**d-f**) Orf8 with a mutation commonly found in the SARS-CoV-2 genome, Orf8-S84L, shows the same effects on histone PTMs H3K9me3 (d), H3K27me3 (e), and H3K9ac (f). N = 332 (GFP), 237 (S84L) cells for H3K9me3; 186, 166 cells for H3K27me3; 332, 237 cells for H3K9ac from 2 independent transfections. (**g**) Orf8 with a 6 amino acid deletion outside of the ARKSAP motif shows the same effects on H3K9me3. N = 216 (GFP), 120 (Orf8), 88 (AGSKSP) cells from 2 independent transfections. n.s., nonsignificant, ***, p<0.001. (b,g) 1-way ANOVA with post-hoc 2-sided t-test and Bonferroni correction. (d-f) 2-sided t-test. Scale bar = 10μM.

**Supplemental Figure 5.**
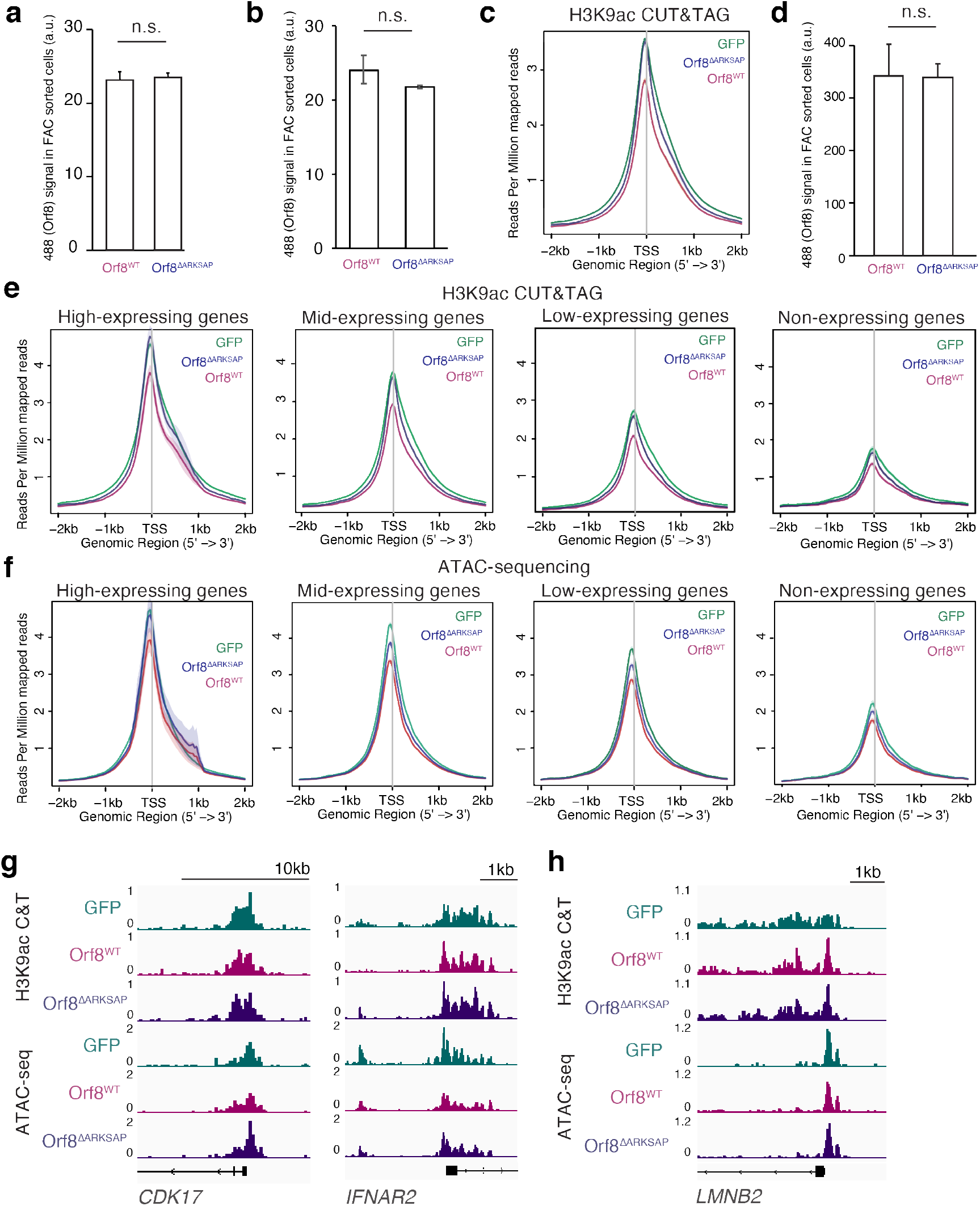
Orf8 effects on chromatin accessibility. (**a**) FAC sorted cells used for western blot analysis express equivalent levels of WT Orf8 and Orf8^ΔARKSAP^. (**b**) FAC sorted cells used for H3K9ac CUT&TAG express equivalent levels of WT Orf8 and Orf8^ΔARKSAP^. (**c**) H3K9ac CUT&TAG sequencing average profile of all expressed genes. (**d**) FAC sorted cells used for ATAC-seq express equivalent levels of WT Orf8 and Orf8^ΔARKSAP^. (**e**) H3K9ac CUT&TAG average profiles for high, mid, low, and non-expressing genes. (**f**) ATAC-seq average profiles for high, mid, low, and non-expressing genes. (**g**) H3K9ac CUT&TAG and ATAC-seq gene tracks of genes relevant to viral responses in HEK cells expression a control plasmid, Orf8, or Orf8^ΔARKSAP^ for genes that show decreased accessibility and H3K9ac (*CDK17* and *INFAR2*). (**h**) While effects of Orf8 are global, genes can be found with only small or no changes with Orf8 in H3K9ac CUT&TAG and ATAC-seq gene tracks. n.s., nonsignificant, (a,c) 2-sided t-test. Error bars (a,c) show standard error.

**Supplemental Figure 6.**
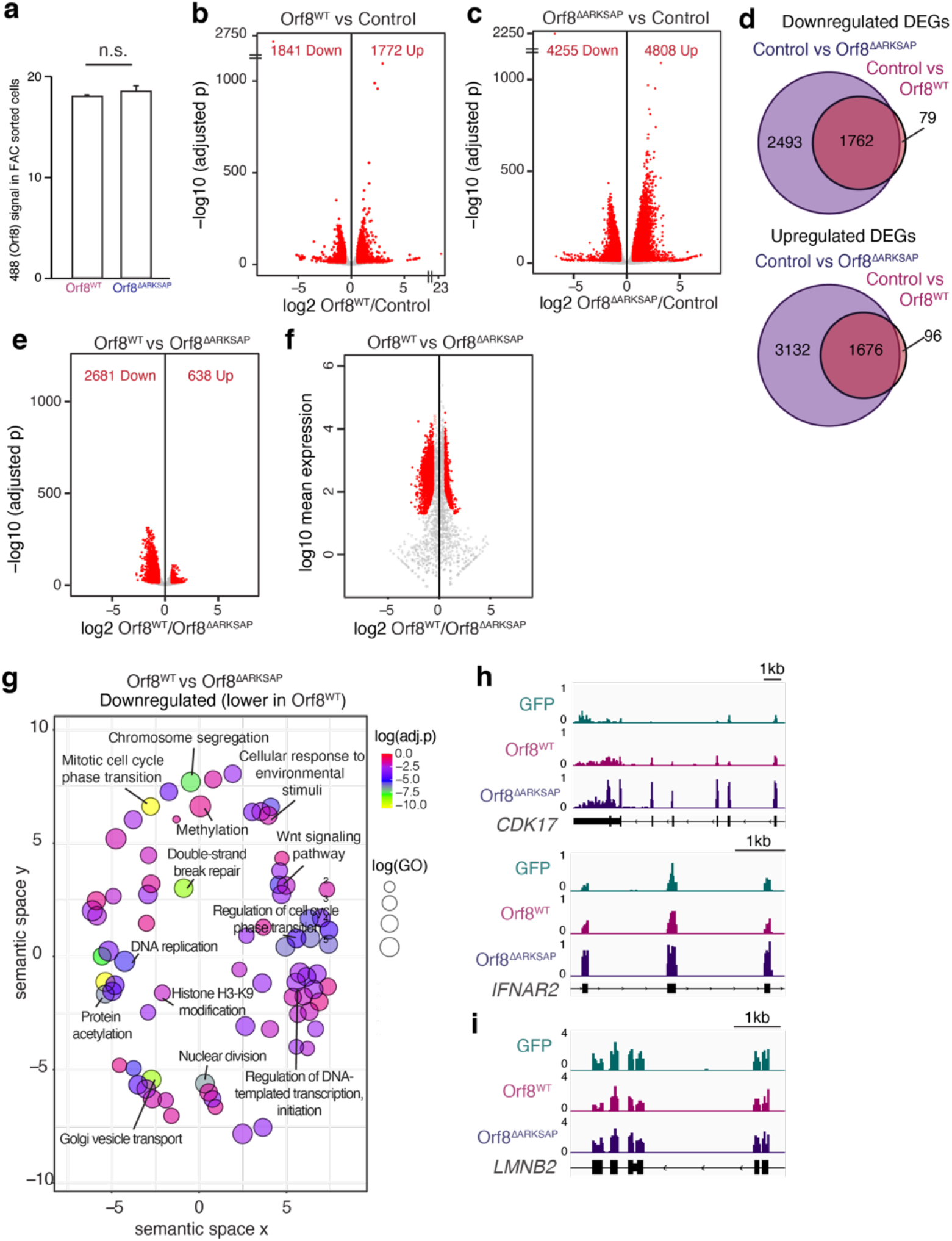
(**a**) FAC sorted cells used for RNA-seq express equivalent levels of Orf8^WT^ and Orf8^ΔARKSAP^. (**b**) Volcano plot of differential gene expression analysis of Orf8^WT^ expressing cells compared to GFP expressing cells. (**c**) Volcano plot of differential gene expression analysis of Orf8^ΔARKSAP^ expressing cells compared to GFP expressing cells. Red indicates significantly differentially expressed genes (adjusted p value < 0.05). (**d**) Overlap of genes down or upregulated by Orf8 and Orf8^ΔARKSAP^ compared to GFP expressing cells. (**e**) Volcano plot of differential gene expression analysis of Orf8^WT^ expressing cells compared to Orf8^ΔARKSAP^ expressing cells. (**f**) Gene ontology analysis of genes that are downregulated by Orf8 compared to Orf8^ΔARKSAP^. (**g**) Gene tracks of genes that are induced by Orf8^ΔARKSAP^ but show a dampened response to Orf8. (**h**) Gene tracks of a gene that is not disrupted by Orf8 expression. n.s., nonsignificant, (a) 2-sided t-test. Error bars (a) show standard error.

**Supplemental Figure 7.**
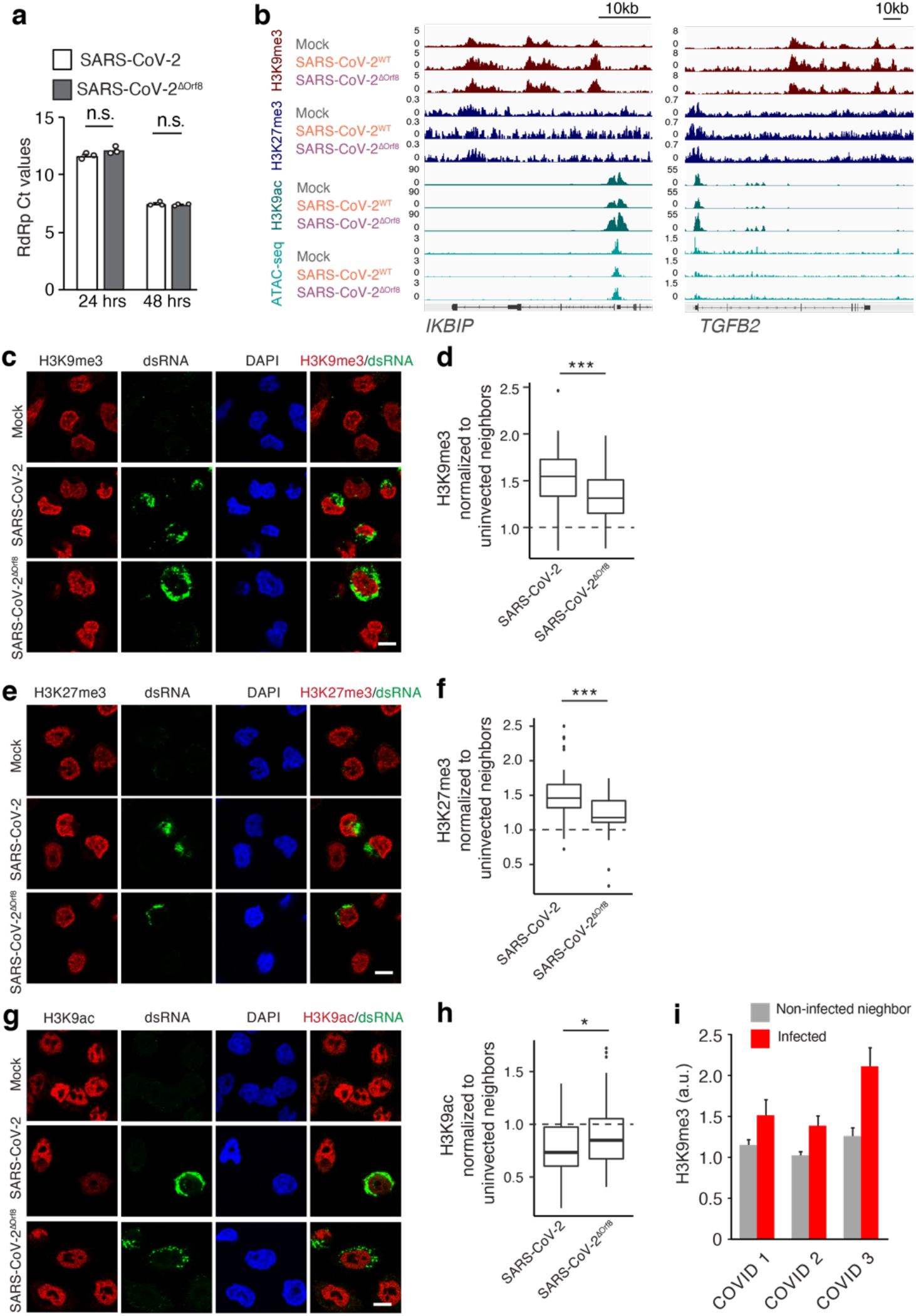
Orf8 mediates SARS-CoV-2 effects on histone PTMs. (**a**) qRT-PCR analysis of expression of SARS-CoV-2 gene RdRp analysis of viral titer in A549^ACE^ pulmonary cells at 24 (a) and 48 (b) hours after infection with SARS-CoV-2^WT^ or SARS-CoV-2^ΔOrf8^ at MOI=1. n = 3 replicates from infection done in parallel. (**b**) ChIP and ATAC-seq gene tracks of genes in signaling pathways relevant to viral response. (**c**,**e**,**g**) H3K9me3 (c), H3K27me3 (e) or H3K9ac (g) staining of A549^ACE^ cells 24 hours after SARS-CoV-2^WT^, SARS-CoV-2^ΔOrf8^, or mock infection at MOI=1. (**d**,**f**,**h**) Quantification of H3K9me3 (d), H3K27me3 (f) or H3K9ac (h). For H3K9me3 n = 39 (SARS-CoV-2), 35 (SARS-CoV-2^ΔOrf8^), for H3K27me3 n = 48 (SARS-CoV-2), 37 (SARS-CoV-2^ΔOrf8^), for H3K9ac n = 94 (SARS-CoV-2), 120 (SARS-CoV-2^ΔOrf8^) cells normalized to uninfected neighbor cells from 1-2 independent infections. Dotted line indicates relative signal in Mock infected condition. (**i**) Quantification of H3K9me3 in infected cells compared to neighboring cells from the same tissue slice for each individual patient sample shown separately. Cell numbers are: sample 1, N=3 infected and 48 uninfected, sample 2, N=7 infected and 125 uninfected, sample 3, N=5 infected and 55 uninfected. *, p<0.05. ***, p<0.001. (d,f) 1-way ANOVA with post-hoc 2-sided t-test and Bonferroni correction. (a-b) 2-sided t-test. Scale bar = 10μM.

**Supplemental Figure 8.**
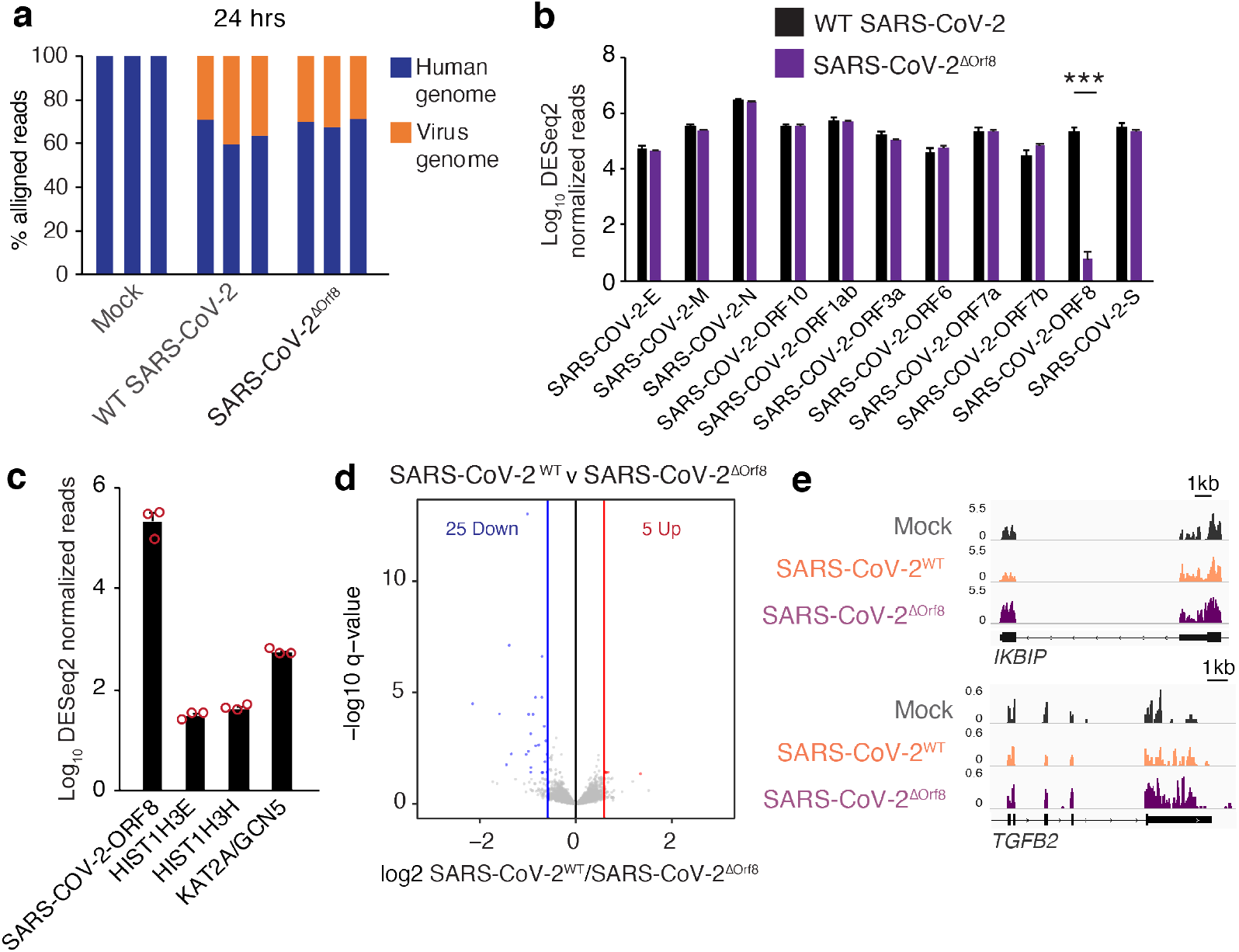
RNA-sequencing of A549^ACE^ cells at 24 hours after infection with SARS-CoV-2^WT^ or SARS-CoV-2^ΔOrf8^. (**a**) Reads aligned to the human and SARS-CoV-2 genomes. (**b**) Levels of SARS-CoV-2 transcripts in cells infected with SARS-CoV-2^WT^ or SARS-CoV-2^ΔOrf8^. (**c**) Normalized reads of SARS-CoV-2 Orf8 and human transcripts of histone H3 genes and KAT2A in cells infected with SARS-CoV-2^WT^. (**d**) Differential gene expression between cells infected with SARS-CoV-2^WT^ and SARS-CoV-2^ΔOrf8^. (**e**) Gene tracks of RNA-seq of cells infected with SARS-CoV-2^WT^ or SARS-CoV-2^ΔOrf8^. ***, p<0.001. 2-way ANOVA with post-hoc 2-sided t-test and Bonferroni correction.

**Supplemental Figure 9.**
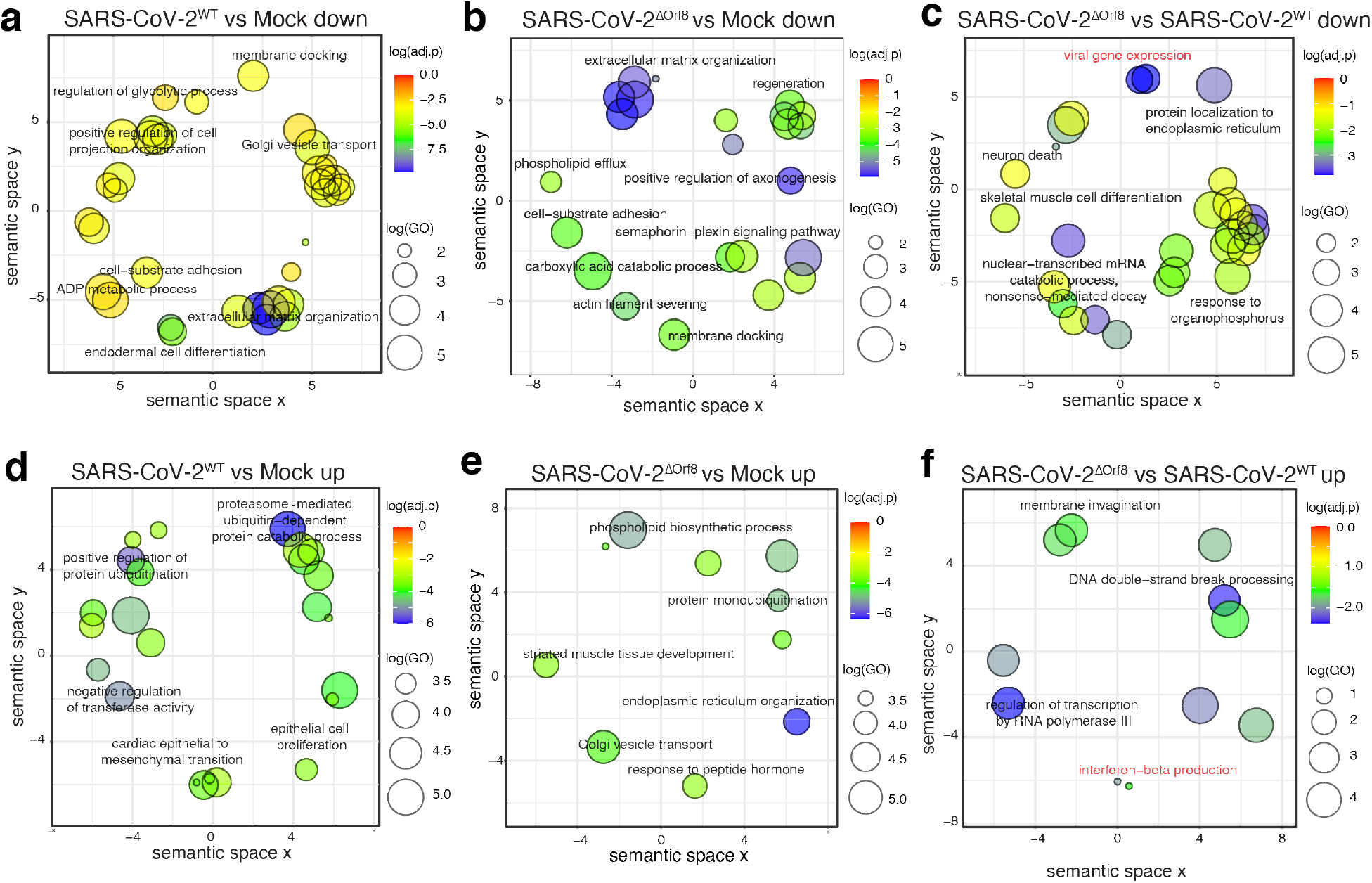
Gene ontology analysis of differentially expressed genes in A549^ACE^ cells at 24 hours after infection with SARS-CoV-2^WT^ or SARS-CoV-2^ΔOrf8^. (**a**) GO analysis of genes downregulated in SARS-CoV-2^WT^ compared to mock infection. (**b**) GO analysis of genes downregulated in SARS-CoV-2^ΔOrf8^ compared to mock infection. (**c**) Gene groups downregulated in SARS-CoV-2^ΔOrf8^ compared to SARS-CoV-2^WT^. (**d**) GO analysis of genes upregulated in SARS-CoV-2^WT^ compared to mock infection. (**e**) GO analysis of genes upregulated in SARS-CoV-2^ΔOrf8^ compared to mock infection. (**f**) GO analysis of genes upregulated in SARS-CoV-2^ΔOrf8^ compared to SARS-CoV-2^WT^. All plots show Revigo clustering of all significantly enriched biological process gene ontology terms with representative terms labeled for each cluster. Red text indicates terms indicating differences in viral response.

**Supplemental Figure 10.**
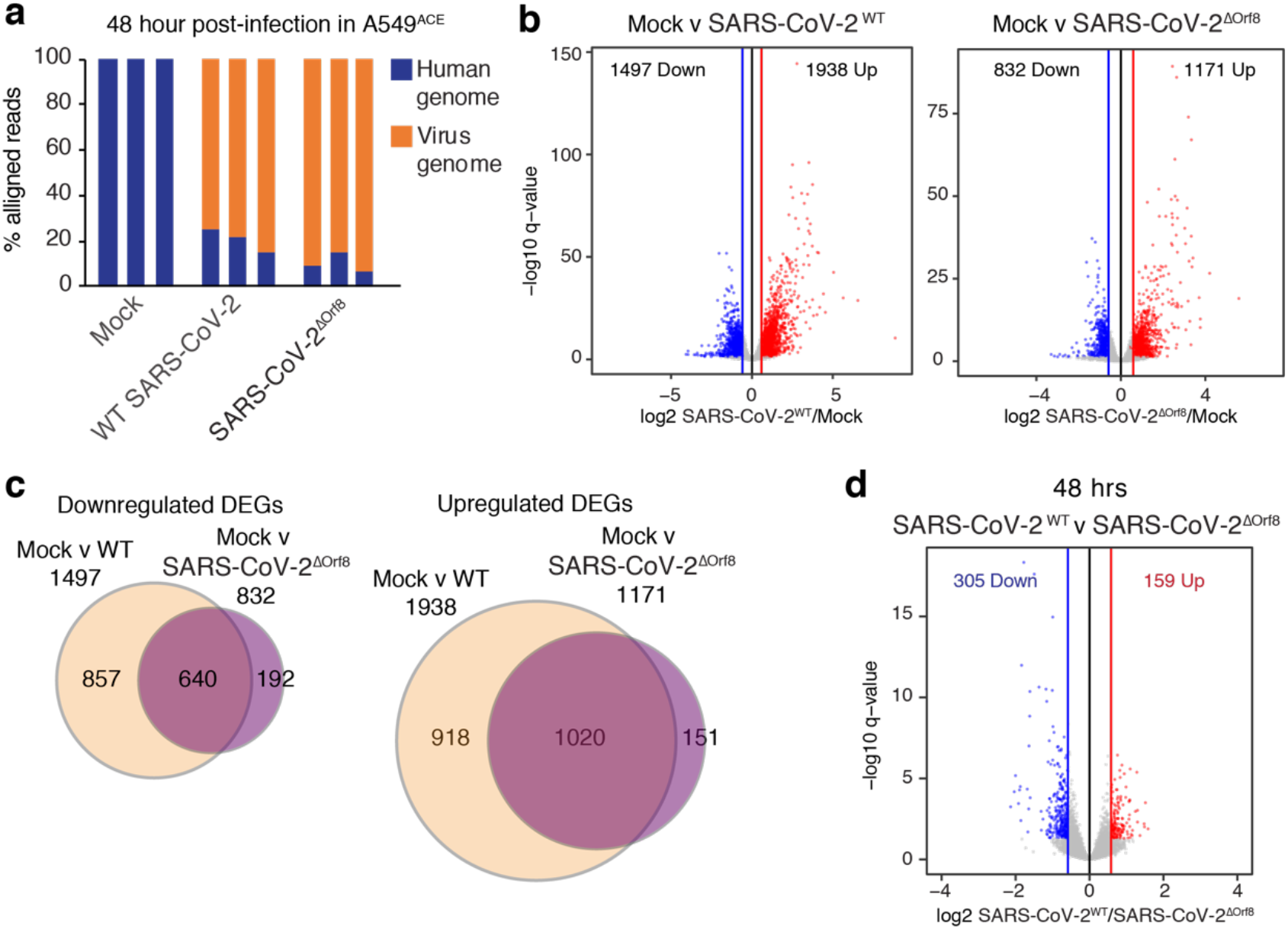
RNA-sequencing of A549^ACE^ cells at 48 hours after infection with SARS-CoV-2^WT^ or SARS-CoV-2^ΔOrf8^. (**a**) Reads aligned to the human and SARS-CoV-2 genomes. (**b**) Differential gene expression analysis of RNA-seq of A549^ACE^ cells 48 hours after SARS-CoV-2^WT^, SARS-CoV-2^ΔOrf8^, or mock infection at MOI=1. Significantly differentially expressed genes (relative to mock infection) are shown in blue (down) and red (up). (**c**) Overlap of differentially expressed genes in response to SARS-CoV-2^WT^ and SARS-CoV-2^ΔOrf8^ infection. (**d**) Differential gene expression between cells infected with SARS-CoV-2^WT^ and SARS-CoV-2ΔOrf8. N = 3

**Supplemental Figure 11.**
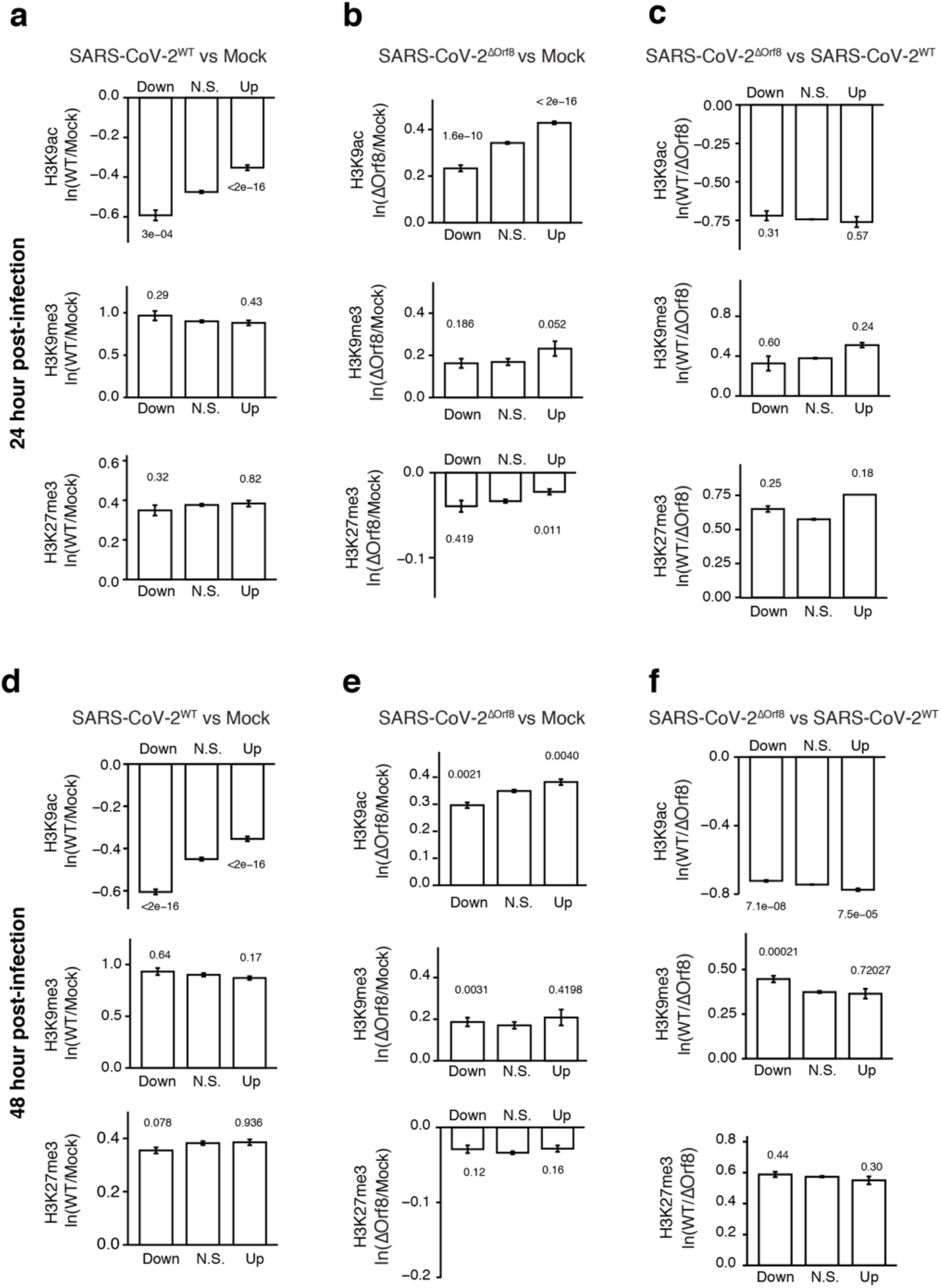
Comparison of ChIP-sequencing and RNA-sequencing data in A549^ACE^ cells. (**a**) Mean peak for SARS-CoV-2^WT^ verses Mock infection for H3K9ac, H3K9me3, or H3K27me3 in genes that are down, unchanged, or upregulated in responses to SARS-CoV-2^WT^ infection compared to Mock infection of A549^ACE^ cells at 24 hours. (**b**) Mean peak for SARS-CoV-2^ΔOrf8^ verses Mock infection for H3K9ac, H3K9me3, or H3K27me3 in genes that are down, unchanged, or upregulated in responses to SARS-CoV-2^ΔOrf8^ infection compared to Mock infection of A549^ACE^ cells at 24 hours. (**c**) Mean peak for SARS-CoV-2^WT^ verses SARS-CoV-2^ΔOrf8^ infection for H3K9ac, H3K9me3, or H3K27me3 in genes that are down, unchanged, or upregulated in responses to SARS-CoV-2^ΔOrf8^ infection compared to SARS-CoV-2^WT^ infection of A549^ACE^ cells at 24 hours. (**d**) Mean peak for SARS-CoV-2^WT^ verses Mock infection for H3K9ac, H3K9me3, or H3K27me3 in genes that are down, unchanged, or upregulated in responses to SARS-CoV-2^WT^ infection compared to Mock infection of A549^ACE^ cells at 48 hours. (**e**) Mean peak for SARS-CoV-2^ΔOrf8^ verses Mock infection for H3K9ac, H3K9me3, or H3K27me3 in genes that are down, unchanged, or upregulated in responses to SARS-CoV-2^ΔOrf8^ infection compared to Mock infection of A549^ACE^ cells at 48 hours. (**f**) Mean peak for SARS-CoV-2^WT^ verses SARS-CoV-2^ΔOrf8^ infection for H3K9ac, H3K9me3, or H3K27me3 in genes that are down, unchanged, or upregulated in responses to SARS-CoV-2^ΔOrf8^ infection compared to SARS-CoV-2^WT^ infection of A549^ACE^ cells at 48 hours. Peak values determined by diffbind for the most significant peak intersecting the TSS of a gene. N.S. indicates nonsignificant genes in DESeq2 analysis. N = 3 ChIP-sequencing for each modification performed in parallel. P values indicate 1-way ANOVA with post-hoc 2-sided t-test and Bonferroni correction.

**Supplemental Figure 12.**
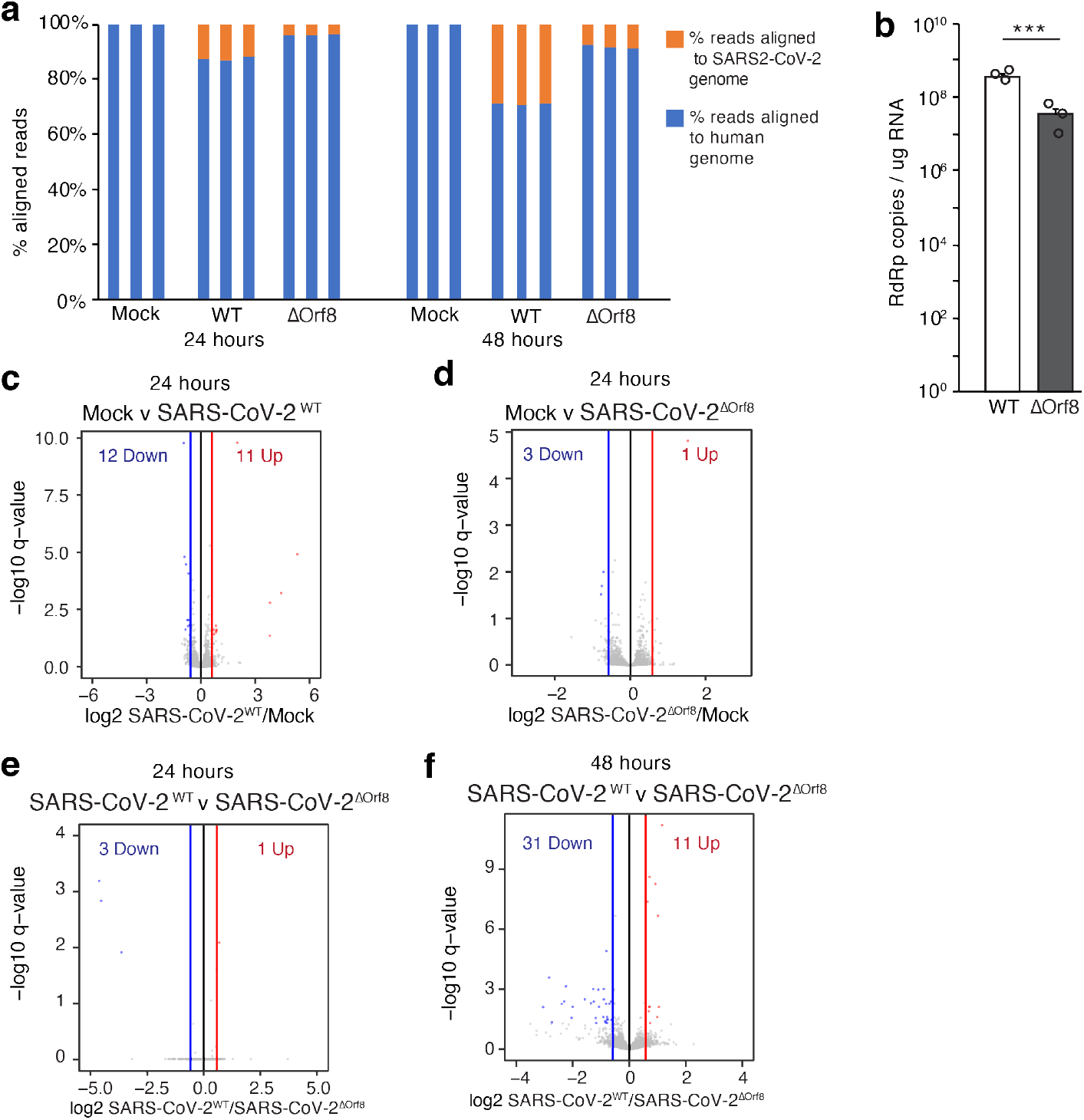
RNA-sequencing of iAT2 cells at after infection with SARS-CoV-2^WT^ or SARS-CoV-2^ΔOrf8^. (**a**) Reads aligned to the human and SARS-CoV-2 genomes at 24 and 48 hours after infection. (**b**) qRT-PCR analysis of expression of SARS-CoV-2 gene RdRp in induced pluripotent stem cell-derived lung alveolar type 2 (iAT2) pulmonary cells at 48 hours after infection with SARS-CoV-2^WT^ or SARS-CoV-2^ΔOrf8^ at MOI=1. N=3 independent replicates performed from a separate derivation of iAT2 cells and separate infection from data shown in Fig. 4. ***, p<0.001, 2-sided t-test. (**c-e**) Differential gene expression analysis of RNA-seq of iAT2 cells 24 hours infection at MOI=1 comparing SARS-CoV-2^WT^ verse mock infection (c) SARS-CoV-2^ΔOrf8^ verse mock (d) and SARS-CoV-2^WT^ verse SARS-CoV-2^ΔOrf8^ (e). (**f**) Differential gene expression analysis of RNA-seq of iAT2 cells 24 hours infection at MOI=1 comparing SARS-CoV-2^WT^ and SARS-CoV-2^ΔOrf8^. Significantly differentially expressed genes (relative to mock infection) are shown in blue (down) and red (up). N = 3.

## METHODS

### A549^ACE2^ cells

ACE expressing A549 cells were generated as previously described^1^. A549-ACE cells were grown in RPMI1640 media with 10% FBS and 1% Pen/Strep and maintained free of mycoplasma. Cells were infected at an MOI of 1 and fixed or lysed at 48 hours after infection.

### HEK cells

HEK293T cells were cultured in DMEM (with 4.5 g/L glucose, L-glutamine and sodium pyruvate), 10% fetal bovine serum (Sigma Aldrich F2442-500ML), and 1% Penicillin-Streptomycin (Gibco 15140122) and maintained free of mycoplasma. Calcium phosphate transfection was used to introduce GFP, Orf8, and Orf8 mutant plasmid DNA to HEK293T cells. Cells were washed 24 hours post-transfection with culture medium and fixed or pelleted and flash frozen 48 hours post-transfection. Cells were fixed using 4% PFA in PBS for 8 minutes. To pellet, cells were detached from the culture plate using TrypLE Express (Gibco 12605010) dissociation reagent, spun down for 5 minutes at 180xg, and flash frozen in liquid nitrogen.

### iAT2 cells

Generation of human-derived induced alveolar epithelial type II-like cells (iAT2) was performed as described^**2**^. To maintain a stable and pure culture of the iAT2 cell line, SFTPC^tdTomato+^ cells were sorted and serially passage every 14 days. Cells were grown as an organoid format using 90% Matrigel with a cell density of 400 cells/ µl. Cells were fed using CK+DCI medium + Rock inhibitor for the first 48h after splitting and then change to K+DCI medium for 5 days followed by CK+DCI medium for 7 days. Every 14 days alveolosphere organoids were passaged, organoids were released from Matrigel using 2mg/ml dispase for 1h at 37°C, then the generation of single cells was reached using 0.05% Trypsin for 15min at 37°C. Cell quantification and viability were assessed using Trypan blue, finally, cells were passaged to new Matrigel drops let them polymerized for 30min at 37°C on a 5% CO2 incubator, after Matrigel solidified cells were fed according to plate format.

For the generation of 2D alveolar cells for virus infection, when alveolospheres organoids were passaged, cells were plated on pre-coated 1/30 matrigel plates at a cell density of 125000 cells/cm2 using CK+DCI medium + Rock inhibitor for the first 48h, and then the medium was changed to CK+DCI medium. 72 hours after cell plating, cells were infected using the SARS-CoV2 virus using MOI:5 for 48h.

### Orf8 constructs

The Orf8 expression plasmid was obtained from Addgene, pLVX-EF1alpha-SARS-CoV-2-orf8-2xStrep-IRES-Puro (Addgene plasmid #141390). Orf8 deletion constructs were produced on the Orf8 backbone using Pfu Turbo HotStart DNA polymerase (Agilent 600322-51) and primers were created using the DNA-based primer design feature of the online PrimerX tool. Constructs were verified by Sanger sequencing.

### SARS-CoV-2 infections

SARS-CoV-2 (USA-WA1/2020 strain) was obtained from BEI and propagated in Vero-E6 cells. The genome RNA was sequenced and found to be identical to GenBank: MN985325.1. Cells were infected with SARS-CoV-2 at MOI=1PFU/cell (A549^ACE2^ and iCM) or MOI=5PFU/cell (iAT2). Virus was added to cells for one hour at 37C, virus removed and replaced with medium. Cells were lysed at 48 hours post infection and RNA isolated. All infections and virus manipulations were conducted in a biosafety level 3 (BSL3) laboratory using appropriate protective equipment and protocols.

### Cell fractionation

Pelleted cells were briefly thawed on ice. Buffer 1 (15mM Tris-HCl (pH 7.5), 60 mM KCl, 15 mM NaCl, 5 mM MgCl2, 1 mM CaCl2, 0.25 M Sucrose with 1 mM PMSF, 1 mM DTT, and Complete Protease Inhibitor cocktail tablet added immediately before use) was added to the pellet at roughly 5 times the volume of the pellet and gently pipetted up and down to dissociate pellet. Samples were incubated on ice for 5 minutes and then an equal volume of buffer 1 with 0.4% NP-40 was added to the sample. Samples were then mixed by inversion for 5 minutes at 4 degrees C. Samples were spun at 200xG for 10 minutes in a prechilled centrifuge to pellet nuclei. The supernatant (cytoplasmic fraction) was transferred to a new tube. Pellets were resuspended gently in 0.5mL buffer 1 to wash nuclei, then pelleted again and supernatant was discarded. Nuclear pellet solubilization buffer (150 mM NaCl, 50 mM Tris-HCl pH 8.0, 1% NP-40, 5 mM MgCl2, with 1 mM PMSF, 1 mM DTT, and Benzonase enzyme at 250U/uL added shortly before use) was added to the pellet at half the volume of buffer 1 used. Samples were then incubated at room temperature in a thermoshaker until the pellet was fully dissolved. Benzonase enzyme was doubled in samples with undissolved material left after 20 minutes. Samples were then centrifuged at 13,000 RPM for 20 minutes at 4 degrees C. Supernatent (nuclei fraction) was collected. Sample concentrations were determined by BCA assay and samples were boiled in a western loading buffer for 10 minutes before analysis by western blot.

### Chromatin sequential salt extraction

Salt extractions were performed as described^3^. Briefly, a 2X RIPA solution was made (100 mM Tris pH 8.0, 2% NP-40, and 0.5% sodium deoxycholate) and mixed with varying concentrations of a 5 M NaCl solution to generate RIPA containing 0, 100, 200, 300, 400, and 500 mM NaCL. pelleted cells were resuspended in buffer A with protease inhibitors (0.3 M sucrose, 60 mM KCl, 60 mM Tris pH 8.0, 2 mM EDTA, and 0.5% NP-40) and rotated at 4 degrees C for 10 minutes. Nuclei were pelleted by centrifugation at 6000xG for 5 minutes at 4 degrees C. Supernatant was removed and saved and 200uL of RIPA with 0mM NaCl and protease inhibitors was added to the sample. Samples were mixed by pipetting 15 times and incubated on ice for 3 minutes before centrifuging for 3 minutes at 6500 xG at 4 degrees C. Supernatant was saved and RIPA steps were repeated for all NaCl concentrations. Samples were boiled and sonicated before analyzing by western blot.

### ATAC-sequencing

HEK cells were stained and sorted to isolate transfected cells using the same as described below. Sorted cells were resuspended in cold Lysis Buffer (10 uL per 10,000 cells; 10 mM Tris-Cl pH 7.5, 10 mM NaCl, 3 mM MgCl2, 0.1% v/v NP-40, 0.1% v/v Tween-20, 0.01% v/v Digitonin), washed in Wash Buffer (10 mM Tris-Cl, pH 7.5, 10 mM NaCl, 3 mM MgCl2, 0.1% v/v Tween-20), and Transposition was performed with Tagment DNA TDE1 (Illumina 15027865). Transposition reactions were cleaned with AMPure XP beads (Beckman A63880), and libraries were generated by PCR with NEBNext High Fidelity 2X PCR Master Mix (NEB M0541). Library size was confirmed on a BioAnalyzer prior to sequencing on the NextSeq 550 (40bp read length, paired end).

For ATAC-sequencing analysis, alignments were performed with Bowtie2 (2.1.0)^4^ using the Hg38 genome using a ChIP-seq pipeline (https://github.com/shenlab-sinai/chip-seq_preprocess). Reads were mapped using NGS plot. For HEK cell ATAC-sequencing, high, mid, low and non-expressing genes were defined by DESeq2 normalized basemean values from HEK cell RNA-sequencing data with under 2 basemean as non-expressing genes and the remaining genes binned into 3 groups for low, mid and high expressing genes.

### ChIP-sequencing

For Orf8 ChIP-sequencing, 2 days after transfection, cells were fixed for 5 minutes with 1% PFA in PBS and then the reaction was quenched with 2.5M glycine. Cells were then washed twice and collected in PBS with protease and phosphatase inhibitors and then pelleted at 1200 rpm for 5 minutes. Cells were then rotated in lysis buffer 1 (50mM HEPES-KOH pH 7.5, 140mM NaCl, 1mM EDTA, 10% glycerol, 0.5% NP-40, 0.25% Triton x-100) for 10 minutes at 4°C and spun at 1350g for 5 minutes at 4°C to isolate nuclei. Supernatant was discarded and cells were resuspended in lysis buffer 2 (10mM Tris-HCl pH 8, 200mM NaCl, 1mM EDTA, 0.5mM EGTA) to lyse nuclei. Cells were rotated for 10 minutes at room temperature and were spun again at 1,350g for 5 minutes at 4°C. The supernatant was discarded and the pellet was resuspended in lysis buffer 3 (10mM Tris-HCl pH 8, 100mM NaCl, 1mM EDTA, 0.5mM EGTA 0.1% EDTA, 0.5% N-lauroylsarcosine). Lysates were sonicated on a Covaris sonicator for 40 minutes (200 cycles/burst). Triton was added to reach a final concentration of 1% and lysates were spun for 10 minutes at 20,000g at 4°C. Streptactin magnetic beads (MagStrep type 3 XT beads, iba #2-4090-002) were added to the lysates overnight rotating at 4°C. Beads were then washed with a low salt buffer (0.1%SDS, 1% triton, 2mM EDTA, 20mM TRIS pH 8, 150mM NaCl), a high salt buffer (0.1%SDS, 1% triton, 2mM EDTA, 20mM TRIS pH 8, 500mM NaCl), a Lithium Choride was buffer (150mM LiCl, 1% NP-40, 1% NaDOC, 1mM EDTA, 10mM TRIS pH 8) and then TE with 50mM NaCl. Chromatin was eluted from beads for 30 minutes shaking at room temperature 55uL BXT elution buffer (iba 2-1042-025) followed by the addition of 150uL elution buffer (50mM Tris-HCl pH8.0, 10mM EDTA, 1% SDS) for 30 minutes at 65°C. Samples were removed from beads and crosslinking was reversed by further incubating chromatin overnight at 65°C. RNA was digested with RNAase for 1 hour at 37°C and protein was digested with proteinase K for 30 minutes at 55°C. DNA was then purified with the Zymo PCR purification kit. The Illumina TruSeq ChIP purification kit was used to prepare samples for sequencing on an Illumina NextSeq 500 instrument (42bp read length, paired end).

For Orf8 ChIP-sequencing analysis, alignments were performed with Bowtie2 (2.1.0)^4^ using the Hg38 genome using a ChIP-seq pipeline (https://github.com/shenlab-sinai/chip-seq_preprocess). Orf8 reads were mapped using NGS plot.

For histone PTM ChIP-sequencing, 4-10M cells were resuspended in 1mL of lysis buffer 1 (50mM HEPES-KOH pH 7.5, 140mM NaCl, 1mM EDTA, 10% glycerol, 0.5% NP-40, 0.25% Triton X-100) and rotated at 4C for 10min, followed by centrifugation removal of supernatant. Cells were then re-suspended in 1mL lysis buffer 2 (10mM Tris-HCl pH 8.0, 200mM NaCl, 1mM EDTA, 0.5mM EGTA), rotated for 10min at 4C, followed by centrifugation removal of supernatant. Cells were then resuspended in 1mL of lysis buffer 3 (10mM Tris-HCl pH 8.0, 100mM NaCl, 1mM EDTA, 0.5mM EGTA, 0.1% Na-Deoxycholate, 0.5% N-lauroylsarcosine) and rotated again for 10min at 4C. This was followed by sonication with a Covaris S220 sonicator for 35 minutes (peak incident power: 140; duty factor: 5 %; cycles/burst: 200). This was followed by addition of 110uL Triton X-100, and centrifugation for 15minutes at maximum speed (20kg) at 4C to clear lysate. Lysate chromatin concentration was then equalized according to DNA content (as measured with a qubit flourometer). Following this, 5% of equivalently-treated chromatin from *C. floridanus* pupae were added to all samples according to chromatin concentration, and 50uL of lysate saved as input shearing control. 250uL of equalized lysate were then added to washed, antibody-conjugated protein A/G Dynabeads (2ug antibody conjugated to 15uL/15uL A/G dynabeads, resuspended in 50uL per IP) and IPs were rotated overnight at 4C in a final volume of 300uL.

The following day, IPs were washed 5x in RIPA wash buffer (50mM HEPES-KOH pH 7.5, 500mM LiCl, 1mM EDTA, 1% NP-40, 0.7% Na-Deoxycholate) and once in TE pH 8.0. Washes were followed by two elutions into 75 µl of elution buffer (50 mM Tris-HCl pH 8.0; 10 mM EDTA; 1% SDS) at 65°C for 45 min with shaking (1,100 RPM). DNA was purified via phenol:chloroform:isoamyl alcohol (25:24:1) followed by ethanol precipitation. Pelleted DNA was resuspended in 25 µl TE. Libraries for sequencing were prepared using the NEBNext® Ultra™ II DNA Library Prep Kit for Illumina® (NEB E7645), as described by the manufacturer but using half volumes of all reagents and starting material. For PCR amplification optimal number of PCR cycles was determined using a qPCR side-reaction using 10% of adapter-ligated, size-selected DNA. 7-10 cycles of PCR were used for hPTM libraries and 5 cycles were used for Input controls. Samples were sequenced on a NextSeq 500 instrument (42bp read length, paired end).

For analysis of histone PTM ChIP-sequencing, reads were demultiplexed using bcl2fastq2 (Illumina) with the options “--mask-short-adapter-reads 20 --minimum-trimmed-read-length 20 -- no-lane-splitting --barcode-mismatches 0”. Reads were trimmed using TRIMMOMATIC (Bolger et al., 2014) with the options “ILLUMINACLIP:[adapter.fa]:2:30:10 LEADING:5 TRAILING:5 SLIDINGWINDOW:4:15 MINLEN:15”, and aligned to a hybrid hg38+*C. floridanus* (v7.5, RefSeq) genome assembly using bowtie2 v2.2.6^4^ with the option “--sensitive-local”. Alignments with a mapping quality below 5 (using samtools) and duplicated reads were removed. Peaks were called using macs2^5^ v2.1.1.20160309 with the options “--call-summits –nomodel –B”. Differential ChIP peaks were called using DiffBind^6^ using the options “bFullLibrarySize=FALSE, bSubControl=TRUE, bTagwise=FALSE” for dba.analyze(). For DiffBind testing the DESeq2 algorithm with blocking was used, and ChIP replicate was used as the blocking factor while testing for differences between Mock and infected samples. For ChIP signal tracks individual replicate tracks were produced for RPM and fold-enrichment over input control, merged, and averaged.

In order to account for potential global differences in hPTM abundance that would otherwise be missed by more standard quantile normalization-type approaches, high-quality de-duplicated read counts were produced for both human-mapping and *C. floridanus*-mapping reads, resulting in proportions of reads mapping to exogenous genome for each hPTM. Input controls were also treated in this way to account for potential differences in initial spike-in addition between samples. For each hPTM, the proportion of spike-in reads were normalized by the appropriate input-control value. Because spike-ins should be inversely proportional to target chromatin concentration, a ratio of CoV/Mock values was produced for each hPTM × replicate, and for CoV2 samples resulting signal values were divided by this ratio. This resulted in per-bp signal values adjusted by the degree of global difference in a given hPTM’s level between sample types.

### RNA-sequencing

RNA was extracted using a Qiagen RNA purification kit. Samples were prepared for sequencing using the Illumina TruSeq purification kit and sequenced on an Illumina NextSeq 500 instrument (75bp read length, single read). Library size was confirmed on a BioAnalyzer prior to sequencing on the NextSeq 550 using single-end, 75 cycles). STAR was used to generate bam files for subsequent TDF file generation using IGVtools.

For RNA-sequencing analysis for SARS-CoV-2 infection experiments, a reference genome for alignment was built by concatenating human (GRCh38 assembly) and SARS-CoV-2 (WA-CDC-WA1/2020 MN985325.1 assembly) genome. For RNA-sequencing analysis for HEK-293T cell experiments, GRCh38 assembly was used. For all RNA-sequencing, reads were aligned using STAR (v2.6.1a) with default parameters and only uniquely mapped reads were retained for downstream analysis. Reads were counted towards human genes (GENCODE v35) and SARS-CoV-2 genes(WA-CDC-WA1/2020 MN985325.1 assembly) using featureCounts (v1.6.2). Low count genes were filtered so that only genes with count per million (CPM) greater than 1 in at least 3 samples were used. Data normalization and differential gene expression analysis was performed using DESeq2 R package (v1.26.0). We defined genes as significant using an FDR cutoff of 0.05 and 1.5X fold change. GO enrichment analysis for differentially expressed genes was implemented using clusterProfiler R package (v3.14.3), using human genome annotation record in org.Hs.eg.db R package (v3.10.0) and BH adjusted p-value of 0.05 as the cutoff.

### Immunoprecipitation

#### Anti–Strep tag affinity purification, whole cell lysate

Protein and binding partners were purified with affinity strep tag purification. For Orf8 PTM analysis and mass spec binding partner analysis, whole cell lysates were prepared as described below. Frozen cell pellets were thawed briefy and suspended in lysis buffer [immunoprecipitation (IP) buffer (50 mM tris-HCl, pH 7.5 at 4°C; 150 mM NaCl, 1 mM EDTA, 10mM sodium butyrate) supplemented with 0.5% Nonidet P 40 Substitute (NP-40; Fluka Analytical) and cOmplete mini EDTA-free protease and PhosSTOP phosphatase inhibitor cocktails (Roche)]. Samples were incubated on a tube rotator for 30 min at 4°C. Debris was pelleted by centrifugation at 13,000 × *g*, at 4°C for 15 min. Lysates were then incubated with streptactin magnetic beads (40ul; MagStrep type 3 XT beads, iba #2-4090-002) for 2 hours rotating at 4°C. Beads were washed three times with 1 ml wash buffer (IP buffer supplemented with 0.05% NP-40) and then once with 1 ml IP buffer. Strep-tagged Orf8 complexes were eluted from beads in Buffer BXT (IBA Lifesciences; Cat. # 2-1042-025) shaking at 1100 rpm for 30 min.

#### Anti–Strep tag affinity purification for chromatin binding partners

Cells were then rotated in lysis buffer 1 (50mM HEPES-KOH pH 7.5, 140mM NaCl, 10mM sodium butyrate, 1mM EDTA, 10% glycerol, 0.5% NP-40, 0.25% Triton x-10) supplemented with 0.5% Nonidet P 40 Substitute (NP-40; Fluka Analytical) and cOmplete mini EDTA-free protease and PhosSTOP phosphatase inhibitor cocktails (Roche)) for 10 minutes at 4°C and spun at 1350g for 5 minutes at 4°C to isolate nuclei. Supernatant was discarded and cells were resuspended in lysis buffer 2 (10mM Tris-HCl pH 8, 200mM NaCl, 10mM sodium butyrate, 1mM EDTA, 0.5mM EGTA) to lyse nuclei. Cells were rotated for 10 minutes at room temperature and were spun again at 1,350g for 5 minutes at 4°C. The supernatant was discarded and chromatin pellet was resuspended in lysis buffer 3 (10mM Tris-HCl pH 8, 100mM NaCl, 10mM sodium butyrate, 1mM EDTA, 0.5mM EGTA 0.1% EDTA, 0.5% N-lauroylsarcosine). Lysates were sonicated using a tip sonicator with 3, 5 second bursts, at 70% power with chilling on ice between bursts. After sonication, lysates were brought to concentration of 1% Triton X-10 to disrupt lamina protein interactions. Debris was pelleted by centrifugation at 16000 × g, at 4°C and supernatant was incubated with streptactin magnetic beads (40ul; MagStrep type 3 XT beads, iba #2-4090-002) for 2 hours rotating at 4°C. Beads were washed three times with 1 ml wash buffer (IP buffer supplemented with 0.05% NP-40) and then once with 1 ml IP buffer. Strep-tagged Orf8 complexes were eluted from beads in Buffer BXT (IBA Lifesciences; Cat. # 2-1042-025) shaking at 1100 rpm for 30 min.

#### Reverse Immunoprecipitation

Chromatin pellet lysate was yielded as described above for chromatin protein immunoprecipitation. Lysates were combined with antibody conjugated protein A Dynabeads (15 ug antibody conjugated to 100ul Dynabeads) and rotated overnight at 4°C. The following day, beads were washed three times with 1 ml wash buffer (IP buffer supplemented with 0.05% NP-40) and then once with 1 ml IP buffer. Chromatin protein complexes were eluted from beads in elution buffer (50mM Tris-HCl pH8.0, 10mM EDTA, 1% SDS) for 30 minutes shaking at 65°C.

### Immunocytochemistry

#### Fluorescent Immunocytochemistry of HEK cells and A549-ACE2 cells

Cells were fixed in 4% paraformaldehyde for 10 minutes, washed with PBS. Fixed cells were permeabilized using 0.5% Triton-X in PBS for 20 minutes. The cells were blocked in blocking solution (PBS, 3% BSA, 2% serum, 0.1% Triton-X) for at least 1 hour and stained with designated primary antibody overnight at 4°C. The following day cells coverslips were washed with PBS incubated with secondary antibodies for 1 hour at room temperature. For detection of strep-tagged Orf8, streptactin DY-488 (IBA Lifesciences; Cat. # 2-1562-050; 1:500) was added to secondary antibody solution. Nuclei were stained with DAPI (1:1000 in PBS) for 10 minutes and washed in PBS. Coverslips were mounted onto microscope slides using ProLong Gold antifade reagent (ThermoFisher).

#### Immunohistological staining of patient lung tissue

Formalin-fixed paraffin-embedded slides were obtained from Penn’s Pathology Clinical Service Center. Slides were deparaffinized and rehydrated as follows: 10 min. Xylene x2, 10 min. 100% Ethanol x2, 5 min. 95% Ethanol, 5 min. 70% Ethanol, 5 min. 50% Ethanol, then running distilled water. Then, slides were processed using heat-induced epitope retrieval (HIER). Slides were incubated in hot sodium citrate buffer (10mM Sodium Citrate, 0.05% Tween-20, pH 6.0), placed in a pressure cooker, and heated in a water bath for 25 minutes with high pressure settings. Slides were cooled at room temperature and washed in TBS x2. Membranes were permeabilized TBS 0.4% Triton-X 100 for 20 min. Slides incubated in blocking solution (TBS; 10% goat serum; 1% BSA; 0.025% Triton-X 100) for 2 hours. Slides were incubated in mouse primary antibody solution of anti SARS-CoV-2 nucleocapsid and rabbit anti-H3K9me3 antibody solution overnight at 4°C. The following day, slides were washed with TBS and incubated in secondary antibody solution. Nuclei were stained with DAPI (5ug/ml) in TBS for 10 min and washed with TBS. Coverslips were mounted with ProLong Gold antifade reagent (ThermoFisher).

### Microscopy

Cells were imaged on an upright Leica DM 6000, TCS SP8 laser scanning confocal microscope with 405 nm, 488, 552, and 638 nm lasers. The microscope uses 2 HyD detectors and 3 PMT detectors. Objectives used were a 63x HC PL APO CS2 oil objective with a NA of 1.40. Type F immersion liquid (Leica) was used for oil objectives. Images were 175.91 × 171.91 miocrons, 1024 by 1024 pixels, and 16-bits per pixel. For PTM quantification, HEK cells and human lung tissue were imaged at a single z-plane and A549 cells were imaged with a z-stack through the nucleus.

### Image analysis

Images were analyzed using Image J software. Single z-plane images of HEK cells and human lung tissue, and summed z-stacks through A549 nuclei were used for PTM quantification. ROI of in-focus nuclei were semi-automatically defined using the DAPI channel and the analyze particles functionality with manual corrections. HEK histone PTMs were quantified in transfected and non-transfected neighboring cells using mean gray values. Strep-tagged Orf8 constructs (Streptactin-488) and GFP signal were used to define transfected cell and HEK histone PTMs levels of transfected cells were relativized to histone PTM levels in non-transfected neighbors. Histone PTMs were quantified in A549 and human lung tissue using integrated density values. dsRNA and SARS-CoV-2 nucleocapsid signal were used to define infected A549s and human lung cells, respectively.

### Protein alignment

To identify potential histone mimicry SARS-CoV-2 protein sequences were aligned to human histone protein sequences (H2A, H2B, H3.1,H3.2 H4, H2A.X, H2A.Z, macroH2A, and H3.3) using Multiple Sequence Comparison by Log-Expectation (MUSCLE) with default settings. SARS-CoV2 protein sequences were obtained from protein sequences published from the first Wuhan isolate.^7^

### Fluorescence activated cell sorting

Frozen cell pellets were gently resuspended in 1 mL FACS buffer (Ca^2+^/Mg^2+^-free PBS, 2% BSA), pelleted at 500 × g at 4°C for 5 minutes, and supernatant was removed. The cells were then gently resuspended in 1 mL FACS buffer with a 1:500 dilution of anti-streptactin antibody and rotated at 4°C for 1 hour, protected from light. Cells were then washed twice in 1 mL FACS buffer, resuspended in 1 mL FACS buffer, and filtered through a 35 uM mesh into FACS tubes. An Aria cells sorter was used to sort positive cells.

### Histone extraction

Transfected cells were isolated by FACS as described above. Sorted cells were pelleted, resuspended in 1 mL cold H2SO4, and rotated overnight at 4°C. Following the overnight incubation, cells were pelleted at max speed and the supernatant was transferred to a fresh tube. TCA was added to 25% by volume and the cells were left on ice at 4°C overnight. Cells were again pelleted at max speed and the supernatant was discarded. Pre-chilled acetone was then used to gently wash the pellet twice. Following the second wash, the tubes were left to air dry before being resuspending in water. Samples were then broken up by alternating 10 minutes of sonication and 30 minutes shaking at 50°C until pellets were fully dissolved.

### Mass spectrometry

#### Histone PTM analysis by quantitative mass spectrometry

Purification of histones was validated by SDS-PAGE followed by Coomassie staining demonstrating sufficient enrichment. A BCA (Thermo Fisher) was performed for protein estimation using the manufacturer’s instructions and 20µg of histone were used for chemical derivatization and digestion as described before^8^. Briefly, unmodified lysines were derivatized twice with a 1:3 ratio of acetonitrile and proprionic anhydride. Histones were then digested with trypsin in a 1:20 enzyme:protein ratio at 37°C overnight. Digested histones with newly formed N-termini were derivatized twice as done before. Finally, histones were desalted as described above.

Dried histone peptides were reconstituted in 0.1% formic acid. A synthetic library of 93 heavy labeled and derivatized peptides containing commonly measured histone PTMs [5] was spiked into the endogenous samples to a final concentration of approximately 100ng/µl of endogenous peptides and 10fm/µl of each heavy labeled synthetic analyte. For each analysis, 1µl of sample was injected on column for data-independent analysis (DIA) analyzed on a Q-Exactive HF (Thermo Scientific) attached to an Ulimate 3000 nano UPLC system and Nanospray Flex Ion Source (Thermo Scientific). Using the same column and buffer conditions as described above, peptides were separated on a 63-minute gradient at 400nL/min starting at 4% buffer B and increasing to 32% buffer B over 58 minutes, and then increasing to 98% buffer B over 5 minutes. The column was then washed at 98% buffer B over 5 minutes and equilibrated to 3% buffer B. Data independent acquisition was performed with the following settings. A full MS1 scan from 300 to 950 m/z was acquired with a resolution of 60,000, ACG target of 3e6, and maximum injection time of 55ms. Then a series of 25 MS2 scans were acquired across the same mass range with sequential isolation windows of 24 m/z with a collision energy of 28, a resolution of 30,000, AGC target of 1e6, and maximum injection time of 55ms. Data analysis and manual inspection using the synthetic library as a reference was performed with Skyline (MacCoss Lab). Ratios were generated using R Studio and statistics carried out in excel.

#### Trypsin & chymotrypsin digestion of Orf8 for identification of Orf8 modifications

The gel band containing Orf8 was destained with 50 mM Ammonium bicarbonate with 50% acetonitrile (ACN). The band was then reduced in 10 mM dithiothreitol in 50 mM ammonium bicarbonate for 30 min at 55°C. Next, the band was alkylated with 100 mM iodoacetamide in 50 mM ammonium bicarbonate at RT for 30 min in the dark. The protein was then digested by incubation with chymotrypsin or trypsin in approximately 1:20 enzyme:protein ratio at 37°C overnight. Following digestion, the supernatant was collected. To extract additional peptides from the gel, 150 μL of 50% ACN and 1% trifluoroacetic acid (TFA) was added and incubated with constant shaking for 30min. The supernatant was collected and 100 μL of ACN was added and incubated with constant shaking for 10min. The final supernatant was collected. All three supernatants were combined and dried. The dried samples were reconstituted by 0.1% TFA and desalted with the C18 micro spin column (Harvard Apparatus). The column was prepared with 200 μL of 100% ACN and equilibrated with 200 μL of loading buffer 0.1% TFA. Peptides were loaded onto the column, washed with a loading buffer, and eluted with 200 μL of 70% acetonitrile in 0.1% formic acid (FA). All steps for loading, washing, and elution were carried out with benchtop centrifugation (300 × g for 2 min). The eluted peptides were dried in a centrifugal vacuum concentrator.

#### Orf8 vs control IP for identification of binding partners

Orf8 immunoprecipitation elutants were reduced and alkylated as described above. Proteins were then digested and desalted with mini S-Trap (Protifi LLC) using the manufacturer’s instructions. Briefly, 25 μL of elutant was combined with 25 μL of 10% sodium dodecyl sulfate (SDS) to a final SDS concentration of 5% SDS after alkylation. Samples were then acidified with phosphoric acid and precipitated by adding 90% methanol (MeOH) in 100 mM triethylammonium bicarbonate (TEAB) in a 6:1 (volume:volume) ratio. Protein was then added to the trap with benchtop centrifugation (4,000 × g for 1 min), washed, and digested with trypsin in a 1:10 enzyme:protein ratio at 37°C overnight. Following digestion, peptides were eluted from the trap with 40 μL of 100 mM TEAB, 40 μL 0f 0.2% FA, and 40 μL of 50% ACN in 0.2% FA. Combined elutant volumes were then dried.

#### Chymotrypsin LC-MS/MS and LC-PRM/MS analysis

Dried peptides were reconstituted with 0.1% FA, and 2 µg of each sample was injected. Chymotrypsin digested Orf8 samples were analyzed on a Q-Exactive (Thermo Scientific) coupled to an Easy nLC 1000 UHPLC system and Nanospray Flex Ion Source (Thermo Scientific). The LC was equipped with a 75 µm × 20 cm column packed in house using Reprosil-Pur C18 AQ (2.4 µm; Dr. Maisch GmbH, Germany). Using aqueous solution of 0.1% FA as buffer A and organic solution of 80% ACN 0.1% FA as buffer B, peptides were separated on a 85 minute gradient at 400nL/min starting at 3% buffer B and increasing to 32% buffer B over 79 minutes, then increasing to 50% buffer B over 5 minutes, and finally increasing to 90% buffer B over 1 minute. The column was then washed at 90% buffer B over 5 minutes and equilibrated to 3% buffer B. Data dependent acquisition was performed with dynamic exclusion of 40 seconds. A full MS1 scan from 350 to 1200 m/z was acquired with a resolution of 70,000, ACG target of 1e6, and maximum injection time of 50ms. Then, a series of MS2 scans were acquired for the top 15 precursors with a charge state of 2-7, a collision energy of 28, and an isolation window of 2.0 m/z. Each MS2 scan was acquired with a resolution of 17,500, AGC target of 2e5, and maximum injection time of 50ms. A database search was performed using the human SwisProt sequence and Orf8 sequence with Proteome Discoverer 2.4 (Thermo Scientific) with the following search criteria: carboxyamidomethylation at cysteine residues as a fixed modification; oxidation at methionine, acetylation at lysine, mono-, di-, and tri-methylation at lysine residues as variable modifications; two maximum allowed missed cleavage; 10 ppm precursor MS tolerance; a 0.2 Da MS/MS. An unscheduled parallel reaction monitoring method^9^ was developed to identify 45 possible modified and unmodified peptide targets of Orf8. Peptides were separated with the same LC gradient conditions. A full MS1 scan from 300 to 900 m/z was acquired with a resolution of 70,000, ACG target of 1e6, and maximum injection time of 50ms. Then, a series of MS2 scans were acquired with a loop count of 23 precursors, a collision energy of 28, and an isolation window of 1.2 m/z. Each MS2 scan was acquired with a resolution of 17,500, AGC target of 1e6, and maximum injection time of 100ms. Data analysis and manual inspection was performed with Skyline^10^ (MacCoss Lab) and IPSA^11^.

#### Trypsin Orf8 LC-MS/MS and LC-PRM/MS analysis and IP LC-MS/MS analysis

Dried peptides were reconstituted with 0.1% FA, and 2 µg of each sample was injected. Data dependent acquisition runs were analyzed on a Q-Exactive HF or HF-X (Thermo Scientific) attached to an Ulimate 3000 nano UPLC system and Nanospray Flex Ion Source (Thermo Scientific). Using the same column and buffer conditions as described above, peptides were separated on a 112 minute gradient at 400nL/min starting at 5% buffer B, increasing to 35% buffer B over 104 minutes, and then increasing to 60% buffer B over 8 minutes. The column was then washed at 95% buffer B for 5 minutes and equilibrated to 5% buffer B. Data dependent acquisition was performed with dynamic exclusion of 45 seconds. A full MS1 scan from 380 to 1200 m/z was acquired with a resolution of 120,000, ACG target of 3e6, and maximum injection time of 32ms. Then, a series of MS2 scans were acquired for the top 20 precursors with a charge state of 2-5, a collision energy of 28, and an isolation window of 1.2 m/z. Each MS2 scan was acquired with a resolution of 30,000, AGC target of 1e6, and maximum injection time of 32ms (HF) or 55ms (HF-X). A database search was performed using the human SwisProt sequence and Orf8 sequence with Proteome Discoverer 2.3 (Thermo Scientific) with the following search criteria: carboxyamidomethylation at cysteine residues as a fixed modification; oxidation at methionine, acetylation, mono-, di-, and tri-methylation at lysine residues as variable modifications; two maximum allowed missed cleavage; 10 ppm precursor MS1 tolerance; a 0.2 Da MS2 tolerance. An unscheduled parallel reaction monitoring method^9^ was developed to identify 16 possible modified and unmodified peptide targets of Orf8. Peptides were separated with the same LC gradient conditions. A full MS1 scan from 350 to 950 m/z was acquired with a resolution of 120,000, ACG target of 3e6, and maximum injection time of 100ms. Then, a series of MS2 scans were acquired with a loop count of 16 precursors, a collision energy of 28, and an isolation window of 1.2 m/z. Each MS2 scan was acquired with a resolution of 30,000, AGC target of 1e6, and maximum injection time of 100ms. Data analysis and manual inspection was performed with Skyline^10^ (MacCoss Lab) and IPSA^11^.

### Antibodies

| Target | Species | Source | Cat # | Dilution | Assay used |
| --- | --- | --- | --- | --- | --- |
| H3K9me3 | Rb | Active Motif | 39161 | 500 | HEK/A549 ICC, IHC |
| H3K9me3 | Rb | Abcam | Ab8898 | 2ug/rxn | ChIP |
| H3K9ac | Ms | Active Motif | 61251 | 500 | HEK ICC |
| H3K9ac | Rb | Active Motif | 39917 | 500 | A549 ICC, CHIP |
| H3K27me3 | Rb | Active Motif | 39155 | 500 | HEK/A549 ICC, ChIP |
| H3K27ac | Ms | Active Motif | 39134 | 500 | HEK ICC, ChIP |
| Lamin-B1 | Rb | Abcam | Ab16048 | 500 | HEK ICC, IP, WB |
| Lamin A/C | Ms | Santa Cruz | sc-376248 | 200 | HEK ICC |
| Orf8_ab1 | Rb | Bioworld | NCP0036 | 250 | A549 ICC |
| Orf8_ab2 | Rb | Bioworld | NCP0010 | 250 | A549 ICC |
| dsRNA | Ms | Jena Bioscience | RNT-SCI-10010200 | 200 | A549 ICC |
| HP1alpha | Rb | Abcam | Ab109028 | 5000 | IP, WB |
| H3 | Rb | Abcam | Ab1791 | 10,000 | IP, WB |
| KAT2A | Rb | Cell Signaling | Ab3305 | 2000 | IP, WB |
| SP2 | Rb | ThermoFisher | PA5-35984 | 1000 | WB |
| Streptactin DY-488 | NA | IBA Lifesciences | 2-1562-050 | 500 | HEK, A549 ICC, FACS |
| Streptactin HRP | NA | IBA Lifesciences | 2-1502-001 | 5000 | WB |
| SARS-CoV-2 nucleocapsid | Rb | ThermoFisher | MA1-7403 | 15 | IHC |
| Anti-mouse AF488 | Gt | ThermoFisher | A-11001 | 500 | ICC, IHC |
| Anti-rabbit AF647 | Gt | ThermoFisher | A-21244 | 500 | ICC, IHC |
| Anti-mouse AF647 | Dk | ThermoFisher | A-21235 | 500 | HEK ICC |
| Anti-rabbit AF568 | Gt | ThermoFisher | A10042 | 500 | HEK/A549 ICC |

ICC = immunocytochemistry, WB = Western blot, IHC = immunohistochemistry

### Data analysis and availability

Box and whisker plots show center line median, box limits for upper and lower quartiles, whiskers for 1.5x interquartile range, and points are outliers. ANOVA testing and plots were generated with R. Bonforonni corrections were applied for multiple comparisons. Fiji was used for image analysis. Imaging and analysis were performed with experimenter blinded to experimental condition where ever possible. For some instances, such as for patient tissue imaging, analysis required targeted selection, imaging, and analysis of infected cells compared to uninfected cells. This required the experimenter was aware of cell infection status while imaging. However, in these cases, the measurement of interest (such as a histone modification stain) was not viewed prior to choosing fields to avoid biasing selection. All genome-wide sequencing data is available under accession number GSE186628 at https://www.ncbi.nlm.nih.gov/geo/query/acc.cgi?acc=GSE186628.

## References

1. Elde, N. C. & Malik, H. S. The evolutionary conundrum of pathogen mimicry. Nat. Rev. Microbiol. 7, 787–797 (2009).

2. Davey, N. E., Travé, G. & Gibson, T. J. How viruses hijack cell regulation. Trends Biochem. Sci. 36, 159–169 (2011).

3. Chaurushiya, M. S. et al. Viral E3 ubiquitin ligase-mediated degradation of a cellular E3: viral mimicry of a cellular phosphorylation mark targets the RNF8 FHA domain. Mol. Cell 46, 79–90 (2012).

4. Marazzi, I. et al. Suppression of the antiviral response by an influenza histone mimic. Nature 483, 428–433 (2012).

5. Schaefer, U., Ho, J. S. Y., Prinjha, R. K. & Tarakhovsky, A. The ‘histone mimicry’ by pathogens. Cold Spring Harb. Symp. Quant. Biol. 78, 81–90 (2013).

6. Avgousti, D. C. et al. A core viral protein binds host nucleosomes to sequester immune danger signals. Nature 535, 173–177 (2016).

7. Avgousti, D. C. et al. Adenovirus Core Protein VII Downregulates the DNA Damage Response on the Host Genome. J. Virol. 91, (2017).

8. Tarakhovsky, A. & Prinjha, R. K. Drawing on disorder: How viruses use histone mimicry to their advantage. J. Exp. Med. 215, 1777–1787 (2018).

9. Strahl, B. D. & Allis, C. D. The language of covalent histone modifications. Nature 403, 41–45 (2000).

10. Jenuwein, T. & Allis, C. D. Translating the histone code. Science 293, 1074–1080 (2001).

11. Berger, S. L. The complex language of chromatin regulation during transcription. Nature 447, 407–412 (2007).

12. Menachery, V. D. et al. MERS-CoV and H5N1 influenza virus antagonize antigen presentation by altering the epigenetic landscape. Proc. Natl. Acad. Sci. U. S. A. 115, E1012–E1021 (2018).

13. Schäfer, A. & Baric, R. S. Epigenetic Landscape during Coronavirus Infection. Pathog. Basel Switz. 6, (2017).

14. Poppe, M. et al. The NF-κB-dependent and -independent transcriptome and chromatin landscapes of human coronavirus 229E-infected cells. PLoS Pathog. 13, e1006286 (2017).

15. Hadjadj, J. et al. Impaired type I interferon activity and inflammatory responses in severe COVID-19 patients. Science 369, 718–724 (2020).

16. Blanco-Melo, D. et al. Imbalanced Host Response to SARS-CoV-2 Drives Development of COVID-19. Cell (2020) doi:10.1016/j.cell.2020.04.026.

17. Li, Y. et al. SARS-CoV-2 induces double-stranded RNA-mediated innate immune responses in respiratory epithelial-derived cells and cardiomyocytes. Proc. Natl. Acad. Sci. U. S. A. 118, e2022643118 (2021).

18. Chan, J. F.-W. et al. Genomic characterization of the 2019 novel human-pathogenic coronavirus isolated from a patient with atypical pneumonia after visiting Wuhan. Emerg. Microbes Infect. 9, 221–236 (2020).

19. Lu, R. et al. Genomic characterisation and epidemiology of 2019 novel coronavirus: implications for virus origins and receptor binding. Lancet Lond. Engl. 395, 565–574 (2020).

20. Yuen, K.-S., Ye, Z.-W., Fung, S.-Y., Chan, C.-P. & Jin, D.-Y. SARS-CoV-2 and COVID-19: The most important research questions. Cell Biosci. 10, 40 (2020).

21. Sampath, S. C. et al. Methylation of a histone mimic within the histone methyltransferase G9a regulates protein complex assembly. Mol. Cell 27, 596–608 (2007).

22. Flower, T. G. et al. Structure of SARS-CoV-2 ORF8, a rapidly evolving coronavirus protein implicated in immune evasion. BioRxiv Prepr. Serv. Biol. (2020) doi:10.1101/2020.08.27.270637.

23. Muth, D. et al. Attenuation of replication by a 29 nucleotide deletion in SARS-coronavirus acquired during the early stages of human-to-human transmission. Sci. Rep. 8, 15177 (2018).

24. Gordon, D. E. et al. A SARS-CoV-2 protein interaction map reveals targets for drug repurposing. Nature (2020) doi:10.1038/s41586-020-2286-9.

25. Hachim, A. et al. ORF8 and ORF3b antibodies are accurate serological markers of early and late SARS-CoV-2 infection. Nat. Immunol. 21, 1293–1301 (2020).

26. Wang, X. et al. Accurate Diagnosis of COVID-19 by a Novel Immunogenic Secreted SARS-CoV-2 orf8 Protein. mBio 11, e02431–20 (2020).

27. Bojkova, D. et al. Proteomics of SARS-CoV-2-infected host cells reveals therapy targets. Nature 583, 469–472 (2020).

28. Zinzula, L. Lost in deletion: The enigmatic ORF8 protein of SARS-CoV-2. Biochem. Biophys. Res. Commun. (2020) doi:10.1016/j.bbrc.2020.10.045.

29. Pereira, F. Evolutionary dynamics of the SARS-CoV-2 ORF8 accessory gene. Infect. Genet. Evol. J. Mol. Epidemiol. Evol. Genet. Infect. Dis. 85, 104525 (2020).

30. Su, Y. C. F. et al. Discovery and Genomic Characterization of a 382-Nucleotide Deletion in ORF7b and ORF8 during the Early Evolution of SARS-CoV-2. mBio 11, (2020).

31. Yiwen Zhang, Junsong Zhang, Yingshi Chen, Baohong Luo, Yaochang Yuan, Feng Huang, Tao Yang, Fei Yu, Jun Liu, Bingfen Liu, Zheng Song, Jingliang Chen, Ting Pan, Xu Zhang, Yuzhuang Li, Rong Li, Wenjing Huang, Fei Xiao, and Hui Zhang. The ORF8 Protein of SARS-CoV-2 Mediates Immune Evasion through Potentially Downregulating MHC-1. BioRxiv (2020).

32. Geng, H. et al. SARS-CoV-2 ORF8 Forms Intracellular Aggregates and Inhibits IFNγ-Induced Antiviral Gene Expression in Human Lung Epithelial Cells. Front. Immunol. 12, 679482 (2021).

33. Rashid, F., Dzakah, E. E., Wang, H. & Tang, S. The ORF8 protein of SARS-CoV-2 induced endoplasmic reticulum stress and mediated immune evasion by antagonizing production of interferon beta. Virus Res. 296, 198350 (2021).

34. Ohki, S., Imamura, T., Higashimura, Y., Matsumoto, K. & Mori, M. Similarities and differences in the conformational stability and reversibility of ORF8, an accessory protein of SARS-CoV-2, and its L84S variant. Biochem. Biophys. Res. Commun. 563, 92–97 (2021).

35. Rashid, F. et al. Mutations in SARS-CoV-2 ORF8 Altered the Bonding Network With Interferon Regulatory Factor 3 to Evade Host Immune System. Front. Microbiol. 12, 703145 (2021).

36. Xie, X. et al. An Infectious cDNA Clone of SARS-CoV-2. Cell Host Microbe 27, 841-848.e3 (2020).

37. Xie, X. et al. Engineering SARS-CoV-2 using a reverse genetic system. Nat. Protoc. 16, 1761–1784 (2021).

38. Li, Y. et al. SARS-CoV-2 induces double-stranded RNA-mediated innate immune responses in respiratory epithelial derived cells and cardiomyocytes. BioRxiv Prepr. Serv. Biol. (2020) doi:10.1101/2020.09.24.312553.

39. Silvas, J. A. et al. Contribution of SARS-CoV-2 Accessory Proteins to Viral Pathogenicity in K18 Human ACE2 Transgenic Mice. J. Virol. 95, e0040221 (2021).

40. Young, B. E. et al. Effects of a major deletion in the SARS-CoV-2 genome on the severity of infection and the inflammatory response: an observational cohort study. Lancet Lond. Engl. 396, 603–611 (2020).

41. Fong, S.-W. et al. Robust Virus-Specific Adaptive Immunity in COVID-19 Patients with SARS-CoV-2 Δ382 Variant Infection. J. Clin. Immunol. (2021) doi:10.1007/s10875-021-01142-z.

42. Lin, X. et al. ORF8 contributes to cytokine storm during SARS-CoV-2 infection by activating IL-17 pathway. iScience 24, 102293 (2021).

43. Valcarcel, A., Bensussen, A., Álvarez-Buylla, E. R. & Díaz, J. Structural Analysis of SARS-CoV-2 ORF8 Protein: Pathogenic and Therapeutic Implications. Front. Genet. 12, 693227 (2021).

44. Lee, S. et al. Virus-induced senescence is a driver and therapeutic target in COVID-19. Nature (2021) doi:10.1038/s41586-021-03995-1.

45. Wang, R. et al. SARS-CoV-2 Restructures the Host Chromatin Architecture. bioRxiv 2021.07.20.453146 (2021) doi:10.1101/2021.07.20.453146.

## Method References

1. Li, Y. et al. SARS-CoV-2 induces double-stranded RNA-mediated innate immune responses in respiratory epithelial derived cells and cardiomyocytes. BioRxiv.

2. Jacob, A. et al. Derivation of self-renewing lung alveolar epithelial type II cells from human pluripotent stem cells. Nat. Protoc. 14, 3303–3332 (2019).

3. Porter, E. G., Connelly, K. E. & Dykhuizen, E. C. Sequential Salt Extractions for the Analysis of Bulk Chromatin Binding Properties of Chromatin Modifying Complexes. J. Vis. Exp. JoVE (2017) doi:10.3791/55369.

4. Langmead, B. & Salzberg, S. L. Fast gapped-read alignment with Bowtie 2. Nat. Methods 9, 357–359 (2012).

5. Zhang, Y. et al. Model-based analysis of ChIP-Seq (MACS). Genome Biol. 9, R137 (2008).

6. Rory Stark<.Ac.Uk>, G. B. C. DiffBind. (Bioconductor, 2017). doi:10.18129/B9.BIOC.DIFFBIND.

7. Wu, F. et al. A new coronavirus associated with human respiratory disease in China. Nature 579, 265–269 (2020).

8. Sidoli, S., Bhanu, N. V., Karch, K. R., Wang, X. & Garcia, B. A. Complete Workflow for Analysis of Histone Post-translational Modifications Using Bottom-up Mass Spectrometry: From Histone Extraction to Data Analysis. J. Vis. Exp. JoVE (2016) doi:10.3791/54112.

9. Peterson, A. C., Russell, J. D., Bailey, D. J., Westphall, M. S. & Coon, J. J. Parallel reaction monitoring for high resolution and high mass accuracy quantitative, targeted proteomics. Mol. Cell. Proteomics MCP 11, 1475–1488 (2012).

10. MacLean, B. et al. Skyline: an open source document editor for creating and analyzing targeted proteomics experiments. Bioinforma. Oxf. Engl. 26, 966–968 (2010).

11. Brademan, D. R., Riley, N. M., Kwiecien, N. W. & Coon, J. J. Interactive Peptide Spectral Annotator: A Versatile Web-based Tool for Proteomic Applications. Mol. Cell. Proteomics MCP 18, S193–S201 (2019).

